# Evaluation of ExoU inhibitors in a *Pseudomonas aeruginosa* scratch infection assay

**DOI:** 10.1101/2020.06.24.170373

**Authors:** Daniel M. Foulkes, Keri McLean, Joscelyn Harris, Atikah S. Haneef, David G. Fernig, Craig Winstanley, Neil Berry, Stephen B. Kaye

## Abstract

*Pseudomonas aeruginosa* has recently been highlighted by the World Health Organisation (WHO) as a major threat with high priority for the development of new therapies. The type III secretion system of *P. aeruginosa* delivers the toxin ExoU into the cytosol of target host cells, where its plasma membrane directed phospholipase activity induces rapid cell lysis. Therefore, inhibition of the phospholipase activity of ExoU would be an important treatment strategy in *P. aeruginosa* infections. We evaluated a panel of ExoU small molecule inhibitors, previously identified from high throughput cellular based assays, and analysed their inhibition of ExoU phospholipase activity *in vitro*. A corneal epithelial (HCE-T) scratch and infection model using florescence microscopy, and cell viability assays, were used to test the efficacy of compounds to inhibit ExoU from *P. aeruginosa*. Compounds Pseudolipasin A, compound A and compound B were effective at mitigating ExoU mediated cytotoxicity after infection at concentrations as low as 0.5 μM. Importantly, by using the antimicrobials moxifloxacin and tobramycin to control bacterial load, these assays were extended from 6 h to 24 h. *P. aeruginosa* remained cytotoxic to HCE-T cells with moxifloxacin, present at the minimal inhibitory concentration (MIC) for 24 h, but, when used in combination with either PSA, compound A or compound B, partial scratch healing was observed. These results provide evidence that ExoU inhibitors could be used in combination with certain antimicrobials as a novel means to treat clinical infections of ExoU producing *P. aeruginosa*.

## Introduction

*Pseudomonas aeruginosa* is a motile, Gram-negative bacterium that causes a wide range of opportunist infections including ocular, soft tissue, urinary tract and respiratory tract infections [1–5]. It is a major cause of intensive care unit-acquired pneumonia (ICUAP), as well as a known coloniser of patients with cystic fibrosis and those who are immunocompromised [6]. Multidrug-resistant *P. aeruginosa* is a major threat, as recently highlighted when the World Health Organisation (WHO) listed carbapenem-resistant *P. aeruginosa* with highest priority for the development of new antibiotics [7]. It is, therefore, imperative that efforts are made to develop novel treatments to target this pathogen. *P. aeruginosa* accounts for ~25% of cases of bacterial keratitis and it is the most common causative agent of bacterial keratitis associated with contact-lens use. After cataracts, bacterial keratitis is the second-largest cause of legal blindness worldwide [8]. During infection, the pathogen exploits its large genome, encoding complex regulatory networks and a wide range of virulence factors, including exotoxins. *P. aeruginosa* employs a needle like apparatus, called the type 3 secretion system (T3SS) that extends through the cell wall to the outer membrane, which it uses to inject certain exotoxins directly into the target cell cytosol [9, 10]. Of the four effector proteins, ExoS, ExoT, ExoU and ExoY, the most cytotoxic is ExoU. In *P. aeruginosa* infections, where the ExoU gene is expressed, disease severity is increased with poorer clinical outcomes [4, 5, 11, 12]. This is considered to be due to rapid cell lysis, mediated by the phospholipase activity of ExoU, which targets the host cell plasma membrane from the cytosol and cannot be halted before conventional antimicrobials can successfully eliminate the pathogen [13, 14].

ExoU is a 74-kDa soluble protein that possesses an N-terminal bacterial chaperone interacting domain followed by a patatin-like phospholipase (PLP) domain and finally a C-terminus containing a 4-helic bundle, which is employed for its insertion into plasma membranes [15–17]. Although the mechanisms of ExoU activation have yet to be fully explored, it is established that certain eukaryotic host co-factors directly interact with ExoU and are required for the induction of its catalytic phospholipase activity [14]. Thus, ExoU relies on binding to ubiquitin and phosphatidylinositol 4,5-bisphosphate (PIP2) to become fully activated [18, 19]. ExoU phospholipase activity can be detected *in vitro* in the presence of ubiquitin as an activating co-factor [20]. In the presence of PIP2, ExoU forms multimers and its ubiquitin dependent catalytic activity is greatly enhanced [21]. In mammalian cells, binding of PIP2 also serves to localise ExoU to the plasma membrane, via the 4-helical bundle domain, where it oligomerises and its catalytic activity induces cellular lysis [18, 22, 23].

The multiple dynamic conformational changes that occur during ExoU activation may be targeted by small molecules to attenuate ExoU activity in clinical infections [14]. Previous cellular based high throughput screening strategies have yielded certain compounds that are able to protect Chinese hamster ovary (CHO) [24] and yeast [25] cells from ExoU mediated cell lysis. Some of these compounds, such as Pseudolipasin A (PSA) have already been shown to inhibit ExoU phospholipase activity *in vitro* [24]. In this study, we investigated the ability of several small molecules and comparatively analysed their abilities to inhibit recombinant ExoU *in vitro*, recombinant ExoU in transfected HeLa cells and performed experiments to observe whether or not certain compounds have the ability to protect corneal epithelial cells from ExoU, produced by *P. aeruginosa*, in a scratch and infection HCE-T cell model. By employing the use of the antimicrobials moxifloxacin and tobramycin, we aimed to control bacterial load in order to extend our scratch and infection assay for longer treatment time courses. We also assessed the ability of antimicrobials to analyse whether or not they could be used in combination with prospective ExoU inhibitors, to rescue cell viability and allow wound healing, in a 24 hour infection assay. Finally, we used molecular docking simulations, employing the crystal structure of ExoU (3TU3), to visualise how compounds could bind to ExoU and the polar contacts they might impart.

## Materials and methods

### Chemicals, reagents and antibodies

Tetracycline (TET), MG132, PMSF, tobramycin and moxifloxacin,α-FLAG and α-tubulin antibodies, Pseudolipasin A (PSA), 5,5’-dithiobis-(2-nitrobenzoic acid) (DNTB) and bovine ubiquitin were purchased from Sigma-Aldrich. The pOPIN bacterial expression vectors were purchased from Addgene. Compounds A (2-[(3-chlorophenyl)amino]-4,6-dimethylnicotinamide) and B 2-[(2,5-dichlorophenyl)amino]-4,6-dimethylnicotinamide were purchased from ChemBridge. Arylsulfonamide 1 was purchased from MolPort. Quinacrine dihydrochloride (QD) and oleyoxylethyl phosphorylcholine (OP) were purchased from Santa Cruz Biotechnology. Phosphatidylinositol 4 5-bisphosphate (PIP2) was purchased from Avanti polar lipids.

### *Pseudomonas aeruginosa* strains and mutants used in this study

The *P. aeruginosa* strains and mutants used in this study have been described previously [15, 26, 27], and were a kind gift from Professor Dara Frank. The ExoU producing strain of *P. aeruginosa*, PA103, and an effector null mutant, which lacks both ExoU and ExoT (PA103ΔUT) were used as positive and negative controls. The PA103ΔUT mutant, when complemented with a pUCP18 plasmid containing the ExoU gene (PA103ΔUT: WT ExoU), fully restores cytotoxic activity towards eukaryotic cells. When the PA103ΔUT mutant is transformed with pUCP18 plasmid encoding the catalytically inactive, S142A ExoU variant (PA103ΔUT: S142A ExoU), acute cytotoxicity is not observed after infection of mammalian cells. *P. aeruginosa* transformed with pUCP plasmids were grown on agar or in LB broth supplemented with 300 μg/ml carbenicillin.

### Recombinant protein production

Full-length ExoU was cloned into pOPINF to generate a His-ExoU encoding construct. Expression conditions were adapted and optimised from previously established protocols [21, 28, 29]. C43(DE3) bacteria were grown in 1 liter of Terrific broth (Melford) supplemented with ampicillin (100 μg/ml) and grown to an optical density (OD_600_) of 0.8 at 30°C before induction of ExoU expression with 0.4 mM isopropyl-β-d-thiogalactopyranoside (IPTG). Phenylmethylsulfonyl fluoride (PMSF) (100 μM) (0.1 % v/v ethanol) was also added to *Escherichia coli* each hour over 3 hours of induced ExoU expression. The bacterial lysis buffer contained 20 mM Tris-HCl pH 8.2, 300 mM NaCl, 0.1 % (v/v) Triton-X-100, 10 mM imidazole, 1 mM DTT, 10 % (v/v) glycerol and a cOmplete protease inhibitor cocktail tablet (Roche) and 100 μM PMSF. ExoU was purified by an initial affinity step (immobilized nickel affinity chromatography) followed by size-exclusion chromatography (16/600 Superdex 200 GE healthcare) in 20 mM Tris-HCl pH 8.2, 100 mM NaCl and 10 % (v/v) glycerol. Recombinant ExoU was frozen in liquid nitrogen and stored at −80C.

### Mass spectrometry

After proteins were resolved by SDS-PAGE and stained with Comassie, gel bands were excised and de-stained using alternating solutions of 25 mM ammonium bicarbonate (AmBic) in 2:1 water/ACN and 25 mM AmBic incubated at 37 °C for 15 min each. Bands were incubated in 10 mM dithiothreitol (DTT) for 60 min at 60 °C, DTT discarded and bands incubated in 55 mM iodoacetamide (IAA) for 45 min at room temperature in the dark. After IAA was discarded, plugs were washed twice in 25 mM AmBic, dehydrated using CAN by washing in acetonitrile and left to air dry. Gel pieces were cooled on ice and rehydrated with 100 μ L of 10 ng/μL Trypsin Gold (Promega, UK) for 5 min. Gel pieces were incubated overnight at 37°C and digestion terminated by adding formic acid to a final concentration of 1% (v/v). The solution was removed and retained. Gel plugs were then incubated in 10 % (v/v) formic acid for 45 min and this solution was combined with the previous solution. The final extract was dried in a vacuum centrifuge until nearly dry and re-suspended in 20 μL of 97:3 water:ACN + 0.1% TFA.

Peptides were analysed on the Ultimate 3000 RSLCTM nano-LC (Thermo Scientific, Hemel Hemstead) coupled to a QExactiveTM mass spectrometer (Thermo Scientific). Peptides from bands 1,2 and 5 were diluted 10-fold in 97:3 water:ACN + 0.1 % (v/v) TFA and 1 μL of this dilution and 1μL of the undiluted peptides from bands 3 and 4 were injected onto the trapping column (Thermo Scientific, PepMap100, C18, 300 μm × 5 mm), using a partial loop injection, for 7 min at a flow rate of 4 μL/min with 0.1% (v/v) FA and then resolved on an analytical column (Easy-Spray C18, 75 μm × 500 mm, 2 μm bead diameter) using a gradient of 97 % A (0.1% v/v) formic acid in H_2_O) and 3 % B (0.1% (v/v) formic acid in 80:20 ACN:H_2_O) to 50% B over 15 min at a flow rate of 300 nl/min. A full-scan mass spectrum was acquired over 350–2200m/z, AGC set to 3e6, with a maximum injection time of 100 ms. The top 3 peaks were selected for MS/MS with an ion selection window of 1.2 m/z and a normalised collision energy of 30, the AGC was set to 1e4 and a maximum injection time of 45 ms. To avoid repeat selection of peptides a 20 sec exclusion window was used.

Data were processed using Proteome Discover 1.4 (Thermo Scientific) and searched using Mascot against an E.coli protein database (retrieved from Uniprot-reviewed proteome accessed May 2015), a database of known contaminates (including common proteases and keratins) and the ExoU protein.

### *In vitro* PLA_2_ assay

The protocol from the Cayman Chemical (USA, Michigan) cPLA2 assay kit was adapted, which allowed the analysis of ExoU sn2 directed phospholipase activity in the presence of compounds in both 96 and 384-well plate formats. Substrate arachidonoyl thio-phosphatidylcholine (ATPC) was purchased from Cayman Chemical as an ethanolic solution. The ethanol was evaporated under a gentle stream of nitrogen gas prior to dissolution of ATPC in 80 mM Hepes pH 7.4, 150 mM NaCl, 4 mM Triton x-100, 30 % (v/v) glycerol and 1 mg/ml bovine serum albumin (BSA) to yield a 1.5 mM substrate stock solution. The final reaction mixture contained 1 μM PIP2, 25 μM ubiquitin, 1 mM ATPC substrate, 2% (v/v) dimethylsulfoxide (DMSO) (with or without compound) and 1.25 mM 5,5-dithio-bis-(2-nitrobenzoic acid) (DTNB) (dissolved in Mili-Q water) to which 1 μM of ExoU was added to initiate substrate hydrolysis. The absorbance at 405 nm (A405) was measured with subtraction of the background absorbance (substrate and DTNB alone) at 10 minute increments over 12 hours. Substrate hydrolysis was calculated using the equation A405/10.00 × 0.05 ml/number of nanomoles of ExoU for the 96-well plate format and A405/10.00 × 0.01 ml/number of nanomoles of ExoU for 384-well plate format, where 10.00 was the path length-adjusted extinction coefficient of DTNB and 0.05 (96-well plate) and 0.01 (384-well plate) were the reaction volumes in milliliters.

### HeLa transfections

Adherent parental Flp-In T-REx-HeLa (Invitrogen) were cultured in Dulbecco’s Modified Eagle medium (DMEM) supplemented with 4 mM L-glutamine, 10 % (v/v) Foetal Bovine Serum (FBS), Penicillin and Streptomycin (Gibco), 4 μg/mL of Blasticidin (Melford) and Zeocin 50 μg/mL (Invitrogen). HeLa cells (2.2 x 10^6^ cells in 10cm dishes and 0.5 x 10^6^ cells for 6-well plates) were seeded 24 hours prior to transfection. For transient transfections, pcDNA5/FRT/TO plasmid encoding Flag-tagged WT ExoU or S142A ExoU was incubated in serum free medium containing lipofectamine 2000 (Invitrogen) for 30 minutes at ambient temperature. The DNA lipofectamine mixture was then added to 10 cm dishes (for Western blot) or 6 well plates (for LDH, microscopy, trypan blue and propidium iodide uptake) of HeLa cells for 12 hours. The cells were then washed with PBS and fresh media containing 1 μg/mL tetracycline (TET) (to induce ExoU expression) and DMSO (0.1% v/v) or indicated compound for indicated time points.

### LDH assays

Lactate dehydrogenase (LDH) release was measured using the Pierce LDH Cytotoxicity Assay Kit (Thermo Scientific) according to the manufactures instructions. For HeLa cell experiments, culture media (50 μL) was assayed at the indicated time points after transfection and induction of ExoU expression with TET, in the presence of indicated compound (0.1% v/v DMSO). The absorbance of the negative controls (untransfected cells) were subtracted to yield the final absorbance values. Medium (50 μL) of HCE-T cells was assayed at time points indicated after infection with the indicated strains of *P. aeruginosa* in the presence of indicated compound (0.1% DMSO, v/v). The results were reported as percent cell lysis normalised to a positive control (according to the manufacturer’s instructions), which gave the maximum amount of observable cell lysis in an appropriate detectable range of absorbance.

### Trypan blue assay

All transfected HeLa cells were collected, after 8 hours of TET induced ExoU expression, by first collecting the culture media (to obtain suspended cells) followed by trypsinisation to procure adherent HeLa cells. The resulting HeLa cell and medium suspension was mixed in a 1:1 ratio with trypan blue reagent (Thermo Scientific) and a Countess II Automated Cell Counter (Thermo Scientific) was employed to detect the percentage of cells with compromised membrane integrity.

### Propidium iodide uptake

Transfected HeLa cells (including suspended cells) were collected by trypsinisation 8 hours after induction of ExoU expression in presence of indicated compound. Samples were stained with propidium iodide (PI) and diluted with PBS so that samples contained less than 500 cells/μl for 10,000 events per run. After gating to select whole cells, employing a BD Accuri C6 flow cytometer, the total cell numbers were evaluated with forward scatter and PI-stained cells were detected by using an appropriate laser for fluorescence with results given as relative florescence units for PI uptake.

### Western blotting

HeLa whole-cell lysates were generated using a modified RIPA buffer (50 mM Tris-HCl pH 7.4, 1% (v/v) NP-40, 0.1% (v/v) SDS, 100 mM NaCl, 1 mM DTT, 10 % (v/v) glycerol, a cOmplete protease inhibitor cocktail tablet (Roche) and 100 μM PMSF. After resuspension in lysis buffer, HeLa cells were briefly sonicated and centrifuged at 16,000 **g** prior to quantification of protein concentration with the Bradford assay. Samples were boiled for 5 minutes in sample buffer (50 mM Ttris-Cl pH 6.8, 1% (w/v) SDS, 10% (v/v) glycerol, 0.01% (w/v) bromophenol blue, and 10 mM DTT). Subsequently, 40 μg of total protein for each sample was resolved by SDS-PAGE prior to transfer to nitrocellulose membranes (Bio-Rad). Membranes were blocked in Tris-buffered saline with+ 0.1% (v/v) Tween 20 (TBS-T) in 5 % (w/v) non-fat dried milk (pH 7.4) followed by incubation with primary and secondary antibodies. Proteins were detected using HRP-conjugated antibodies and enhanced chemiluminesence reagent (Bio-Rad). Band intensities were quantified using ImageJ software.

### HCE-T scratch and infection assay

HCE-T cells were seeded (0.5 x 10^6^) in 6-well plates and grown until a fully confluent monolayer had formed, at which point two parallel scratches were made across the diameter of the wells with a 10 μL pipette tip. In tandem, PA103 and PA103 mutant bacteria were inoculated into LB culture medium (with or without appropriate antibiotic selection marker) and grown overnight. Subsequently, 1 mL of the overnight bacterial culture was subcultured in 50 ml of fresh LB medium and allowed to expand until an OD_600_ of 0.8 was reached. At the point of compound addition to scratched HCT cells, indicated PA103 or PA103 mutant strains were added immediately at a multiplicity of infection (MOI) of 2.5. Where indicated, 2 μM of moxifloxacin or 6 μM of tobramycin was also added to control bacterial growth.

### MTT assays

HCE-T cells were seeded in a 96-well plate at a concentration of 0.2 × 10^6^ cells/ml and allowed to grow until a fully confluent monolayer had formed. Compound addition was performed in triplicate, with all experiments including a final concentration of 0.1% DMSO (v/v). Metabolic activity was quantified 48 hours after compound addition, and cell viability was quantified employing an MTT assay kit (Abcam, place) according to the manufacturer’s instructions. Briefly, thiazolyl blue tetrazolium bromide was dissolved in phosphate buffered saline (PBS) and added to cells at a final concentration of 0.25 mg/ml and incubated at 37°C for 3 hours. The reaction was stopped by the addition of 50 μL acidified 10 % (w/v) SDS, followed by reading of absorbance at 570 nm. Viability was defined relative to DMSO-containing controls incubated for the same time.

### Fluorescence microscopy

HCE-T cells treated as indicated in the figure legends were incubated for either 8 h or 24 h before analysis by florescent microscopy, employing Live/Dead staining (Invitrogen), according to the manufacturer’s instructions, in order to differentiate and visualise viable and dead/dying cells. Briefly, the culture medium was removed from the infected HCT cells in 6 well plates, and washed with 1 ml of PBS three times. Fresh medium containing 5 μM of both calcein (Ex/Em 494/517 nm) and ethidium homodimer-1(Ex/Em 528/617 nm). Images of the scratched HCT cells were obtained on either an Apotome Zeiss Axio Observer or a Nikon Eclipse TiE.

### Molecular docking simulations

The chemical structures of PSA, compound A and compound B were illustrated in ChemDraw and imported into Spartan18 where their 3D structures were energy minimised using MMFF94 forcefield prior to docking experiments. The crystal structure of ExoU (bound to SpcU) (PDB: 3TU3) was imported into the molecular visualising software Hermes and the program GOLD 5.2 [30] was employed to model ligand docking to ExoU with the binding site of the protein defined as being within 10 Å of the catalytic Serine 142 residue of ExoU. Default settings were retained apart from ‘GA settings’ were changed to 200%. Protein and ligand docking poses were visualised in the PyMOL Molecular Graphics System, Version 2.0 Schrödinger. Noncovalent contacts were analysed with ViewContacts software [31].

### Statistical analysis

All experimental procedures were repeated in at least three separate experiments with matched positive and negative controls (unless stated otherwise) and results are presented as means ± SD. When applied, statistical significance of differences (*P ≤ 0.05) was assessed using One-way ANOVA or Student’s t tests for normally distributed data. Statistical tests were performed using either SPSS or Prism 7 (GraphPad Software).

## Results

### Purification and analysis of recombinant ExoU *in vitro*

In order to analyse the *in vitro* phospholipase activity of ExoU, we expressed ExoU with an N-terminal histidine tag in *E. coli* and purified it sequentially by immobilised metal affinity chromatography (IMAC) and then by size-exclusion chromatography (Figure 1A). Building on previously optimised ExoU expression conditions [28, 29], we added 100 μM of the serine protease inhibitor PMSF to *E. coli*, followed by the addition of IPTG to induce expression of ExoU for 3 hours. This afforded enhanced yields of recombinant His-tagged ExoU Supplementary (Figure 1A). Without PMSF, typical yields per litre of *E. coli* culture were 0.1 mg. If however, PMSF was added to IPTG induced *E. coli*, yields of 0.5 mg per litre could be achieved. Mass spectrometry was employed to confirm the presence of ExoU and to elucidate the identity of two contaminants of similar molecular weight. The proteins Glutamine-fructose-6-phosphate aminotransferase (67 kDa) and Bifunctional polymyxin resistance protein ArnA (74 kDa) were only present in *E. coli* after induction of ExoU expression (Figure 1A left). Importantly, despite similar molecular weights, these impurities could be separated from ExoU when it further purified by size-exclusion chromatography (Figure 1A right).

**Figure 1.**
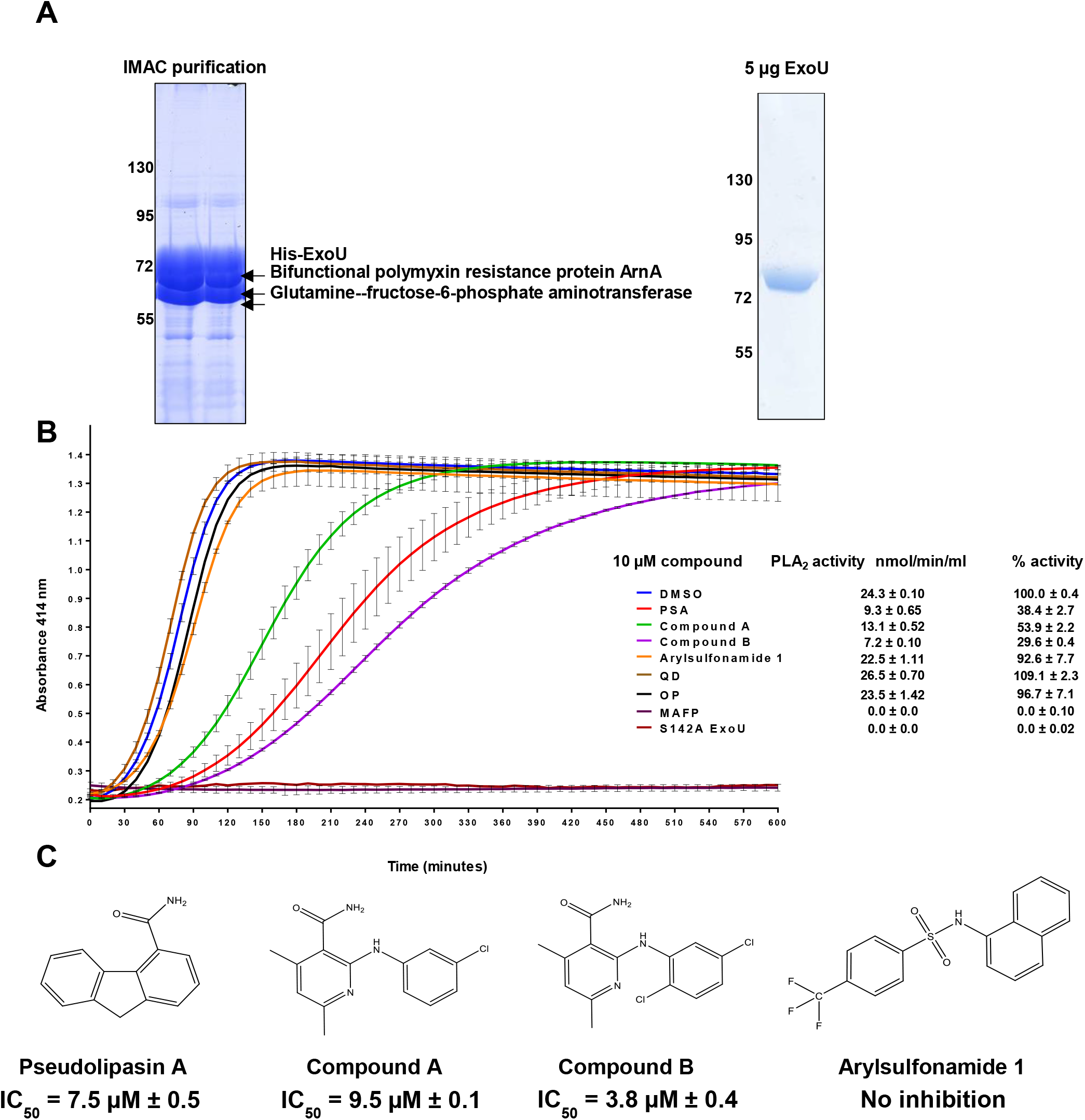
Purification and *in vitro* analysis of recombinant ExoU with prospective small molecule inhibitors. (A) Left: His-tagged ExoU was purified from C43(DE3) *E. coli*. Immobilised metal affinity chromatography (IMAC) purified His-ExoU, including two major contaminant proteins, were resolved by SDS–polyacrylamide gel electrophoresis (SDS-PAGE) and identified by employing mass spectrometry. Right: His-ExoU was further purified to homogeneity by size-exclusion chromatography; 5 μg of His-ExoU was resolved and visualised by SDS-PAGE. (B) The hydrolysis of arachidonoyl Thio-PC substrate by ExoU was assessed in the presence of 10 μM of the indicated compound. To each reaction, ubiquitin and PIP_2_ were added in order to allow induction of ExoU phospholipase activity. Experiments were performed in triplicate, the results represent means, and error bars represent standard deviations and representative profiles are shown of substrate conversion as a function of absorbance with progression of time (QD = quinacrine dihydrochloride and OP = oleyoxylethyl phosphoryl choline). (C) Chemical structures of prospective compounds that are proposed to inhibit ExoU mediated toxicity in cells. IC_50_ values are shown.

The phospholipase activity of ExoU was studied *in vitro* by adaptation of the Cayman-chemical PLA_2_ phospholipase assay kit, so that analysis was compatible with 96 and 384-well plate formats, in the presence of PIP2 and ubiquitin as co-factors for ExoU activation. Arachidonoyl Thio-PC substrate hydrolysis by 1 μM of ExoU in 2% v/v DMSO measured 24.3 ± 0.1 nmol/min/ml (Figure 1B). Serine 142 is indispensable for ExoU catalytic activity [15] and mutation to an alanine abolished detectable substrate hydrolysis (Figure 1B). The promiscuous, broad spectrum PLA_2_ inhibitor MAFP, which covalently binds to the catalytic serine residue of target PLA_2_ enzymes irreversibly inhibited ExoU [24, 32] (Figure 1B). Previously identified ExoU inhibitors Pseudolipasin A (PSA), compound A and compound B [24] (Figure 1C), at a concentration of 10 μM, were inhibitors of ExoU catalytic activity *in vitro*, resulting in decreased rates of substrate hydrolysis to 20 ± 3.8, 40 ± 2.8 and 15 ± 1.1 nmol/min/ml, respectively (Figure 1B). Dose response analysis was performed for compounds PSA, compound A and compound B in order to establish IC_50_ values for inhibition, which were 7.5 ± 0.5, 9.5 ± 0.1 and 3.8 ± 0.4 μM, respectively. The inhibitor arylsulfonamide 1 (Figure 1C) was previously identified from a high throughput screen to protect yeast after infection with *P. aeruginosa* (with ExoU as only virulence effector) [25]. In our analysis, this inhibitor did not inhibit recombinant ExoU phospholipase activity *in vitro*. In addition, we tested two non-specific inhibitors of human PLA_2_ enzymes quinacrine dihydrochloride (QD) and oleyoxylethyl phosphoryl choline (OP), which also did not inhibit *in vitro* ExoU activity (Figure 1B).

### Targeting of ExoU with prospective inhibitors in a HeLa cell model

Flp-In T-REx-HeLa cells were transfected with pcDNA5FRT/TO, encoding either WT or S142A ExoU cDNA, so that FLAG-tagged-ExoU could be expressed upon addition of tetracycline (TET). Transfected HeLa cells induced to express WT ExoU underwent rapid cellular lysis within 8 hours, whereas cells expressing S142A ExoU remained intact (supplementary Figure 2A). This meant that only S142A ExoU could be reasonably observed by immunoblotting analysis (exploiting the N terminal flag tag) of the cell extracts (Figure 2A)[33]. Increasing the amount of transfected WT ExoU cDNA and harvesting cells 4 hours after transfection, allowed the observation of WT ExoU with extended film exposure during immunoblotting (supplementary Figure 3A). An increased amount of FLAG-WT ExoU could be detected when transfected HeLa cells were also treated with 50 μM of the serine protease inhibitor PMSF for 4 hours, but not the proteasome inhibitor MG132 (supplementary Figure 3B).

**Figure 2:**
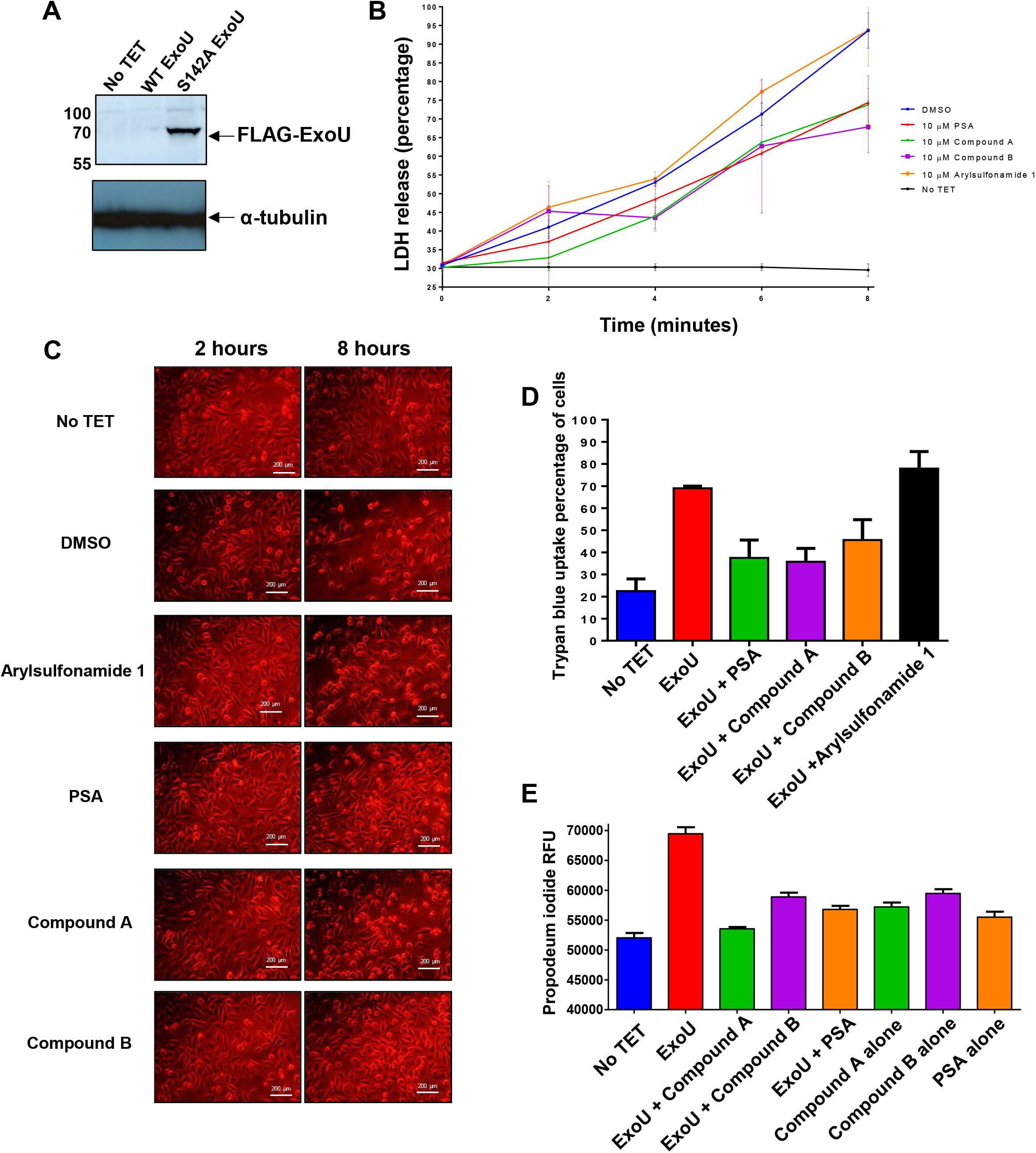
Inhibition of ExoU in a HeLa cell transfection model with prospective small molecules. (A) Flp-In T-REx-HeLa cells were transfected with pcDNA5/FRT/TO encoding the full length WT or S142A EoxU gene so that FLAG-ExoU could be expressed upon incubation with tetracycline (TET) for 6 hours, prior to whole cell lysis and analysis by western blotting. (B) LDH release of HeLa cells transfected and induced to express WT ExoU over 8 hours in the presence of 10 μM of indicated compound. One-way ANOVA analyses were performed to determine statistical significance between DMSO and compound treated cells. (C) Brightfield microscopy images of HeLa cells 8 hours subsequent to induction of WT ExoU epression in the presence of indicated compound. (D) Trypan blue uptake of HeLa cells 8 hours subsequent to induction WT ExoU expression in the presence of indicated compound. (E) Propidium iodide uptake of WT ExoU expressing HeLa cells in the presence of indicated compound, measured by flow cytometry.

In order to assess the efficacy of prospective ExoU inhibitors, transfected HeLa cells were induced to express WT ExoU by the addition of TET for 8 hours in the presence of 10 μM of either PSA, compound A, compound B or arylsulfonamide 1; LDH release was assayed to quantify cell lysis (Figure 2B). With no inhibitor treatment (DMSO-blue) a steady increase in LDH release, up to the maximum detectable activity, was observed over the 8 hour time course. The compound arylsulfonamide 1 (orange) afforded no protection from WT-ExoU mediated cell lysis, whereas PSA (red), compound A (green) and compound B (purple), significantly decreased the quantity of LDH activity detected in the culture medium over 8 hours (p values of 0.01, 0.01 and 0.02, respectively). Brightfield microscopy images (Figure 2C) complemented these findings by revealing observable morphological changes to the transfected HeLa cells when WT-ExoU was expressed. This included cell rounding and membrane blebbing (Figure 2C) [27]. Expression of WT-ExoU, without inhibitor treatment (DMSO), also correlated with fewer cells adhered to the well. Similar cellular morphologies were observed for HeLa cells induced to express ExoU in the presence of 10 μM arylsulfonamide 1, indicating that this compound had no substantial effect on ExoU inhibition. Treatment with either 10 μM PSA, compound A or compound B resulted in more adherent cells with normal morphology and fewer instances of cell rounding and membrane blebbing observed.

Trypan blue and propidium iodide (PI) cellular uptake were employed to further assess cell lysis in transfected HeLa cells, induced to express ExoU after 8 hours in the presence of prospective inhibitors. Consistently, the percentage of cells that absorbed the trypan blue dye was increased when HeLa cells were induced to express ExoU (Figure 2D). Compounds PSA, A and B, but not arylsulfonamide 1 caused a reduction in the uptake of trypan blue in induced HeLa cells. From a baseline uptake of PI, when ExoU expression was not induced (no TET), there was a significant increase PI florescence in cells induced to express ExoU (TET) (Figure 2E). This uptake was abrogated in the presence of either 10 μM PSA, compound A or compound B, but not arylsulfonamide 1.

Western blot analysis was employed to determine the potential effects of compounds on ExoU stability in HeLa cells. As WT-ExoU was not readily detectable in whole cell extracts (Figure 2A), we expressed FLAG-tagged S142A ExoU, which was detected in cellular lysates by immunoblotting, after HeLa cells were TET induced and incubated with chosen compounds for 8 hours. After transfection with pcDNA5/FRT/TO, encoding FLAG-tagged S142A ExoU, and induction with TET for 8 hours in the presence of compound, we observed a decrease in the total abundance of S142A ExoU only in the presence of compounds A and B (Figure 3A). PSA, arylsulfonamide 1 and MG132 did not affect the quantity of S142A ExoU present in cellular lysates in comparison to DMSO. The difference in total S142A ExoU protein, when quantified by densitometry, was significantly less for compound A (p=0.005) and compound B (p=0.008) compared to DMSO (0.1% v/v) alone.

**Figure 3:**
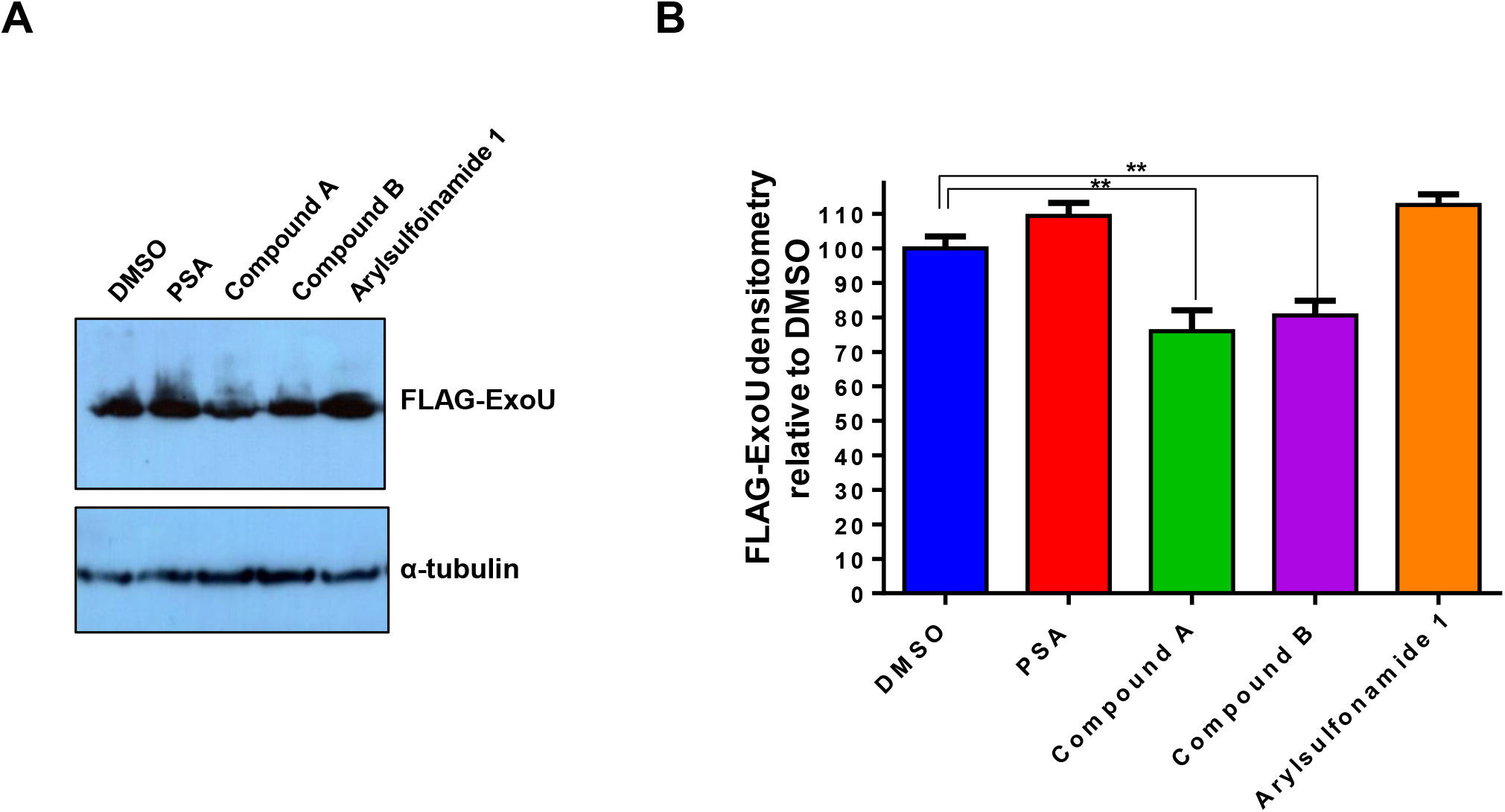
Analysis of S142A ExoU stability in HeLa cells in the presence of compound. (A) Flp-In T-REx-HeLa cells were transfected with pcDNA5/FRT/TO encoding the full length S142A EoxU gene FLAG-S142A ExoU expression was induced with TET and 10 μM of the indicated compound for 8 hours, prior to whole cell lysis and analysis by Western blotting. (B) Densitometry analysis of S142A-ExoU signal from compound treated HeLa cells, relative to a DMSO. T-tests were used to determine statistically significant differences between compound A (p=0.0035) and compound B (p=0.0039) relative to DMSO treated.

### Establishment of a scratch assay to assess toxicity of ExoU expressing *Pseudomonas aeruginosa*

Expanding on established infection models, of eukaryotic cells in culture by ExoU expressing *P. aeruginosa* [26, 27], we developed a human corneal cell (HCE-T) scratch and infection assay. We employed previously utilised bacterial strains to assess the cytotoxic effects of ExoU in the HCE-T cells [26, 27]. The ExoU expressing clinical isolate strain of *P. aeruginosa*, PA103, an ExoU and ExoT knock-out mutant (PA103 ΔUT) complemented with pUCP encoding either WT or S142A ExoU (PA103 ΔUT: ExoU and PA103 ΔUT: S142A ExoU), were used [26]. If either PA103 ΔUT: ExoU (which employs ExoU as the only T3SS cytotoxic effector) was added to fully confluent HCE-T cells, no effects of infection or ExoU mediated cytotoxicity could be detected after 6 hours using fluorescence microscopy and measuring LDH activity (Figure 4A). If, however, a scratch was applied across the diameter of the well prior to infection, lysis of HCE-T cells could by detected and observed by florescence microscopy along the border and at the periphery of the scratch 6 hours after initial infection (Figure 4A). When the scratched HCE-T cells were infected with S142A ExoU expressing PA103, there was a narrowing of the scratch (indicative of wound healing) and no observable dead cells adjacent to the borders (supplementary Figure 4A). This phenomenon was analogous to the control condition of no bacteria added to the scratched HCE-T cells and reflected by the decrease in LDH activity detected relative to PA103 or PA103 ΔUT: WT ExoU infection (supplementary Figure 4B).

**Figure 3:**
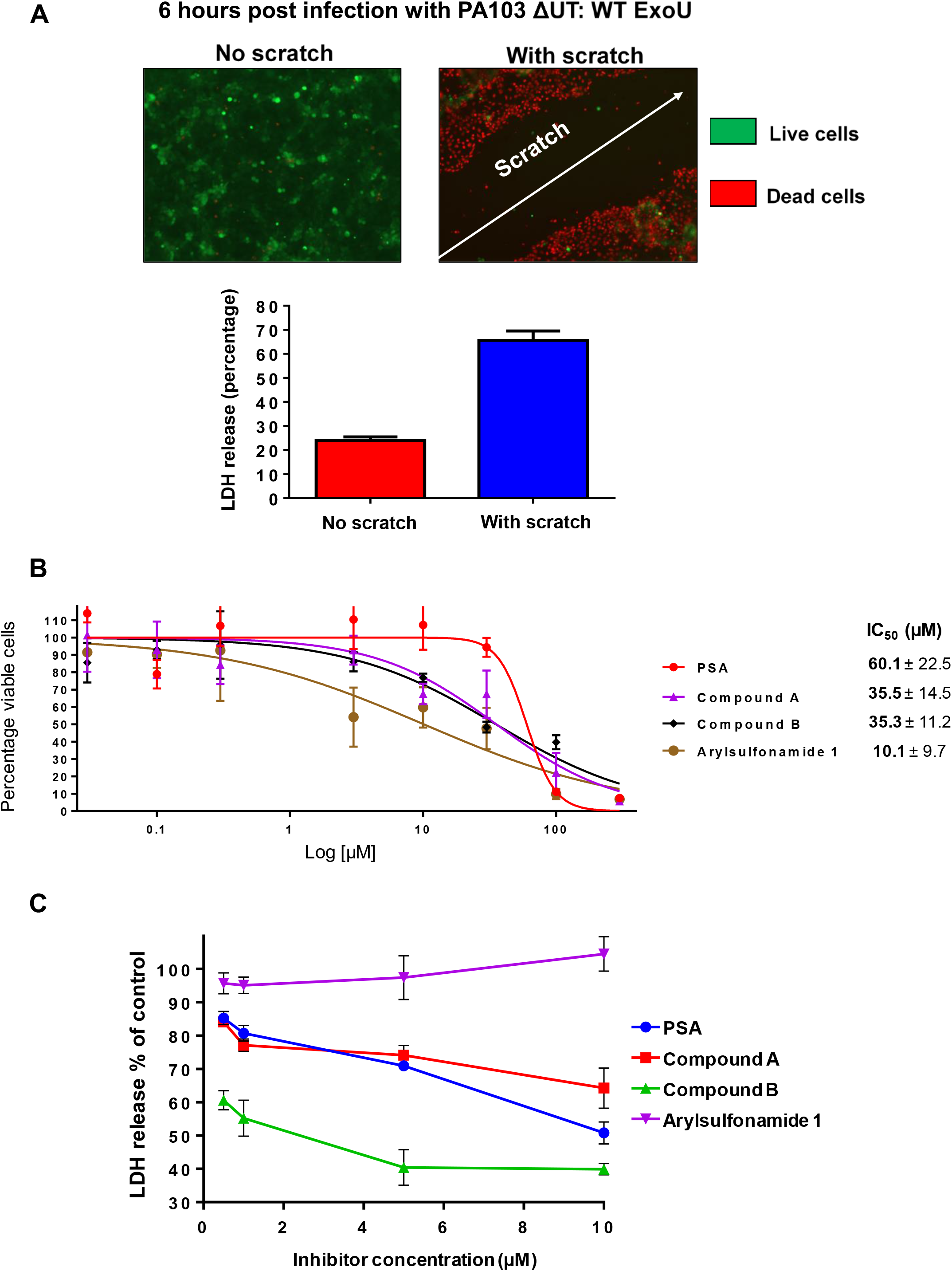
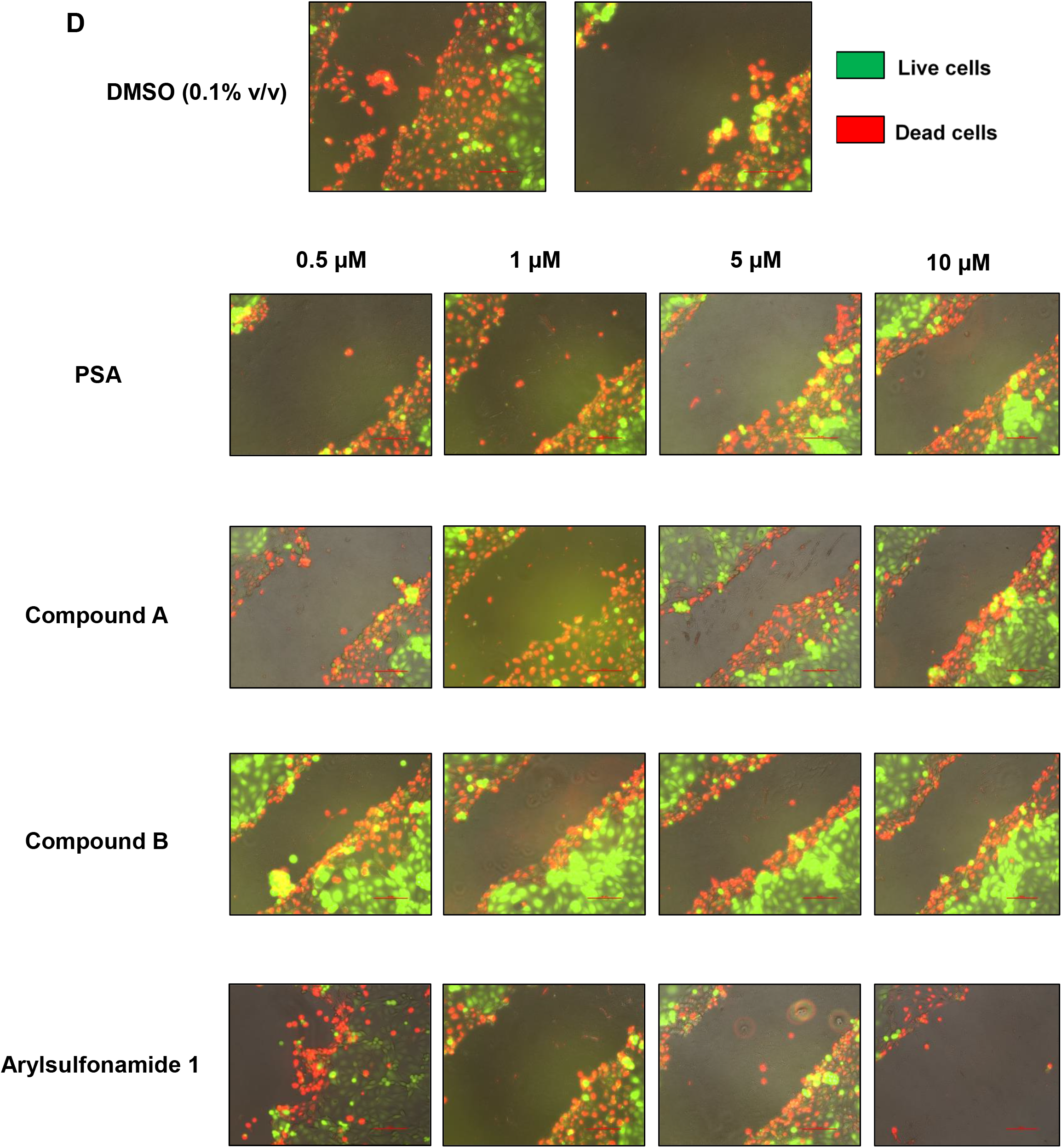
Protection of scratched HCE-T cells, during infection with ExoU expressing PA103, by selected compounds. (A) HCE-T were gown to full confluence and a scratch applied to the cell monolayer prior to infection with PA103 ΔUT: ExoU at an MOI of 2.5 for 6 hours, followed by analysis by Live/Dead fluorescence microscopy (top) or LDH release (bottom). (B) MTT assays comparing the cytotoxicity of ExoU inhibitors in HCE-T cells. The MTT assay was performed 72 hours subsequent to initial compound exposure and IC_50_ values in μM ± SD derived from 3 independent experiments are shown. (C) Dose response analysis of inhibitors analysing protective effect of compounds on scratched then infected HCE-T cells after 6 hours incubation, by LDH release. (D) Live/Dead fluorescence microscopy analysis of scratched HCE-T cells 6 hours post infection, in the presence of varying concentrations of indicated compound.

This scratch and infection assay was used to assess the ability of prospective small molecules to mitigate cytotoxicity induced by ExoU. MTT assays showed that PSA, compound A and compound B were well tolerated, with IC_50_ values of 60.1 ± 22.5, 35.5 ± 14.5 and 35.3 ± 11.2 μM, respectively (Figure 4B). No compound induced toxicity was apparent at assay working concentrations, with the exception of arylsulfonamide 1 (brown), which exhibited an IC_50_ value for the MTT assay of 10.1 ± 9.7 μM (Figure 4B).

Pseudolipasin A, compound A and compound B, all reduced HCE-T LDH release 6 hours after infection with PA103 ΔUT: ExoU, in a dose dependent manner (Figure 4C). Compound B was associated with the greatest reduction in observed LDH release, with decreased LDH activity observed at concentrations as low as 0.5 μM. At 0.5 μM, compound B afforded protection of HCE-T cells from ExoU mediated cell lysis, apparent from 40% reduction in LDH release, compared to no treatment at all (0.1% v/v DMSO control). Indeed, higher concentrations of compound afforded enhanced protection of HCE-T cells; at 10 μM of PSA, compound A and compound B, with an observed reduction in LDH release of 48.8%, 35.4% and 59.7%. Live/dead fluorescence microscopy complemented these data (Figure 4D). Decrease in wound size and fewer dead cells along the border of the scratch was observed for treatments with varying concentrations of PSA, compound A and compound B after infecting scratched HCE-T cells with PA103 ΔUT: ExoU, compared to DMSO (Figure 4D). Consistent with a lack of inhibitory activity on recombinant ExoU phospholipase activity or inhibition of ExoU mediated cytotoxicity in HeLa cells, arylsulfonamide 1 did not protect scratched HCE-T cells against ExoU expressing PA103 infection (Figure 4D).

### Compounds combined with moxifloxacin reduce ExoU induced cytotoxicity over 24 hours

Antimicrobials were employed to control bacterial load and extend the HCE-T scratch/infection assay, allowing observation of compounds’ effects after 24 hours. Without antibiotic present, the bacteria overwhelmed the cell culture and were cytotoxic to HCE-T cells after 24 hours (supplementary Figure 5). Moxifloxacin, at an inhibitory concentration (MIC) of 2 μM (Figure 5A), limited the number of colony forming units (CFU) of bacteria in the culture medium after 24 hours, whilst still enabling T3SS and ExoU mediated cytotoxicity (Figure 5B); extending ExoU induced cytotoxicity and its related inhibition in the scratch assay. In contrast, although tobramycin reduced bacterial growth at the established MIC of 6 μM (Figure 5A), there was no observable ExoU mediated cytotoxicity (Figure 5B). Neither moxifloxacin nor tobramycin inhibited recombinant ExoU phospholipase activity *in vitro* (supplementary Figure 6). Importantly, none of the potential ExoU inhibitors were bactericidal over 24 hours and MICs could not be established, indicating that these inhibitors did not affect bacterial growth.

**Figure 5:**
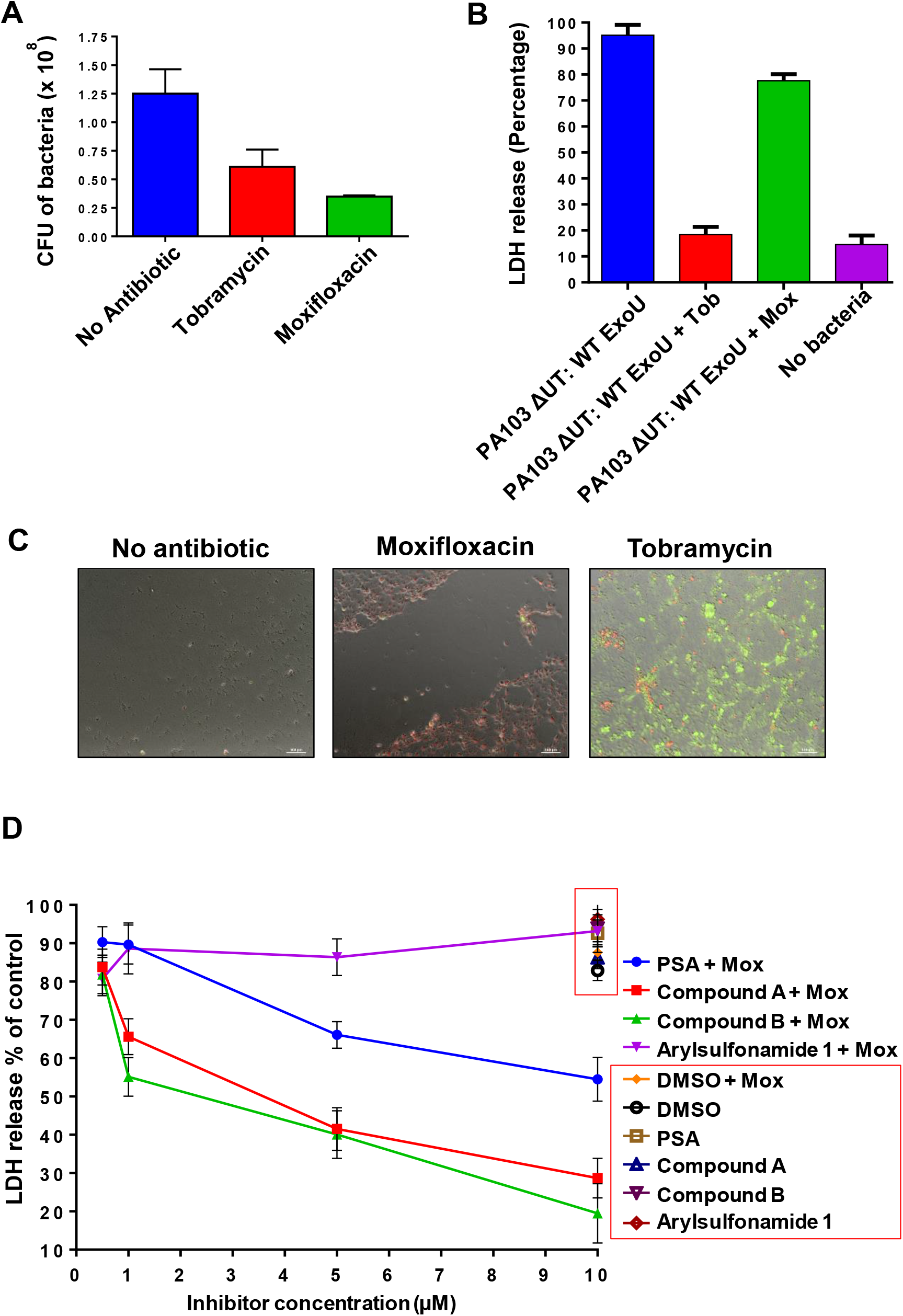
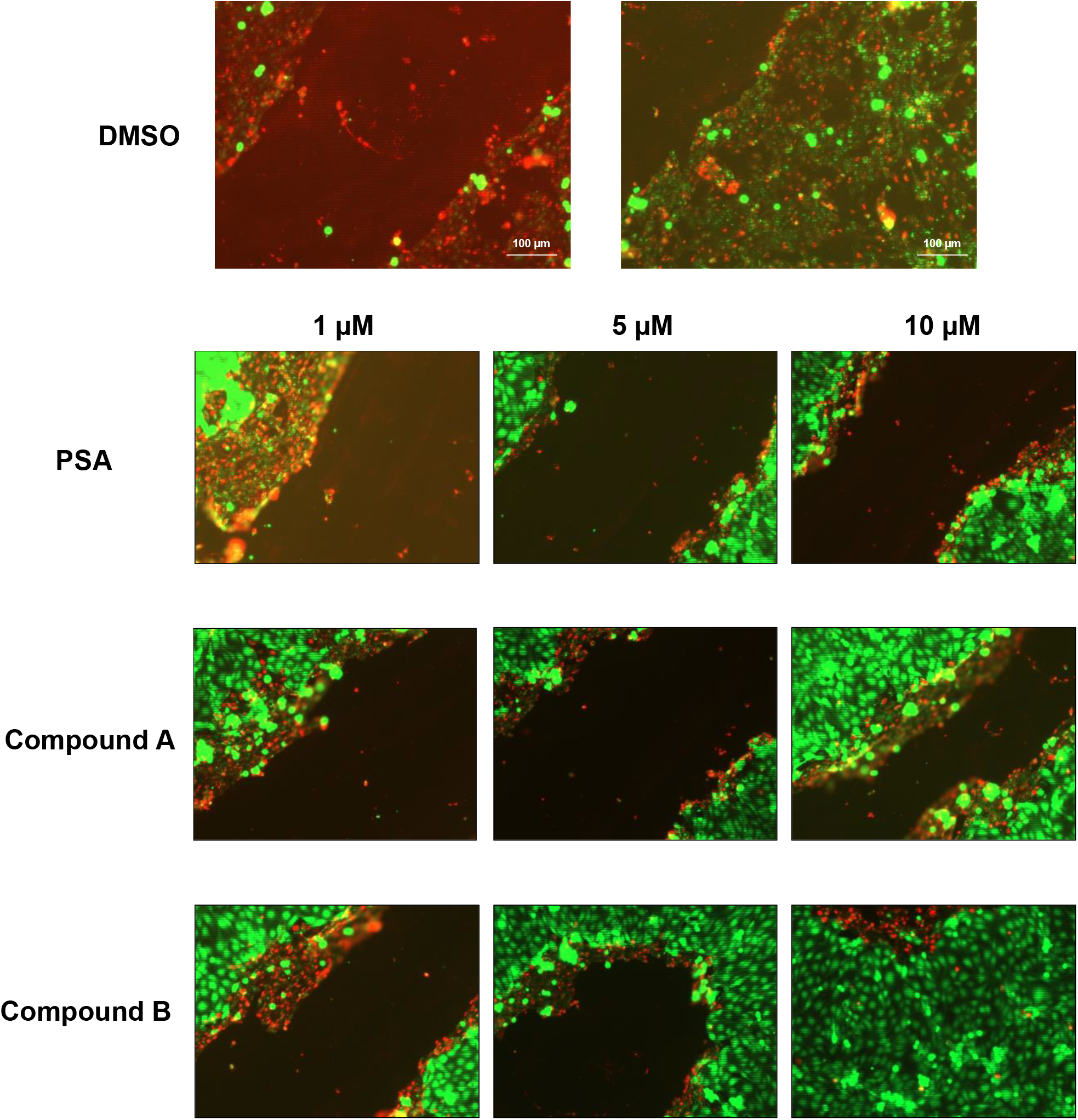
Compounds synergise with moxifloxacin to mitigate cell death induced by ExoU expressing PA103 over 24 hours. (A) Tobramycin (Tob) and moxifloxacin (Mox), at the established MICs of 2 and 6 μM, respectively, were added to scratched HCE-T cells that had been infected with PA103 ΔUT: ExoU at an MOI of 2.5 for 24 hours. The number of colony forming units (CFU) in the cell culture medium was then deduced. (B) LDH release from scratched HCE-T cells after 24 hours infection in the presence of tobramycin or moxifloxacin at MIC. (C) Live/Dead fluorescence microscopy analysis of scratched HCE-T cells 24 hours post infection, without antibiotic present or with tobramycin or moxifloxacin at the MIC. (D) LDH assay for dose response analysis of inhibitors analysing protective effect of compounds on scratched then infected HCE-T cells after 24 hours incubation in the presence or absence of moxifloxacin at the MIC. (E) Live/dead fluorescence microscopy analysis of scratched HCE-T cells 24 hours post infection, in the presence of varying concentrations of indicated compound, with moxifloxacin present at the MIC.

Upon analysis of LDH release 24 hours post infection of scratched HCE-T cells with PA103 ΔUT: ExoU, dose response experiments revealed that PSA, compound A and compound B used in combination with 2 μM moxifloxacin were far more effective at protecting HCE-T cells from lysis than any compound or antibiotic used in isolation (Figure 5D). Combination of compounds A and B with moxifloxacin led to the greatest reduction in cell lysis (Figure 5D), at 66.5% and 75.5% LDH reductions respectively (at a 10 μM concentration).

Live/dead fluorescence microscopy was used to further assess the scratched HCE-T cell assay at 24 hours and complemented LDH release assay data (Figure 5E). For moxifloxacin alone there was extensive ExoU mediated cytotoxicity. Moxifloxacin in combination with PSA, compound A or compound B, was associated with a reduction in scratch width with fewer dead cells along the scratch border. Although the wound did not close completely, for compounds A and compound B there was cell migration and partial wound closure. Strikingly, with tobramycin alone at the MIC, no ExoU cytotoxicity was observed and the wound in the scratched HCE-T cells completely healed.

### Molecular docking simulations of compounds to ExoU

PSA, compound A and compound B displayed characteristics of competitive inhibition of ExoU *in vitro* phospholipase activity (Figure 1B). Using the crystal structure of ExoU, co-crystallised with its cognate chaperone SpcU (PDB: 3TU3) [34], we performed molecular docking simulations to visualise potential compound-ExoU interactions (Figure 6). The Connolly surface of ExoU revealed a potential substrate binding the pocket, adjacent to the catalytic Serine 142 residue (Figure 6A, yellow), as a potential ligand docking site. We observed that certain compounds (PSA in Figure 6B and compound B in Figure 6C) could dock with favourable energetics into this solvent exposed region. All the highest scoring docking solutions revealed that the compounds tested had similar poses.

**Figure 6:**
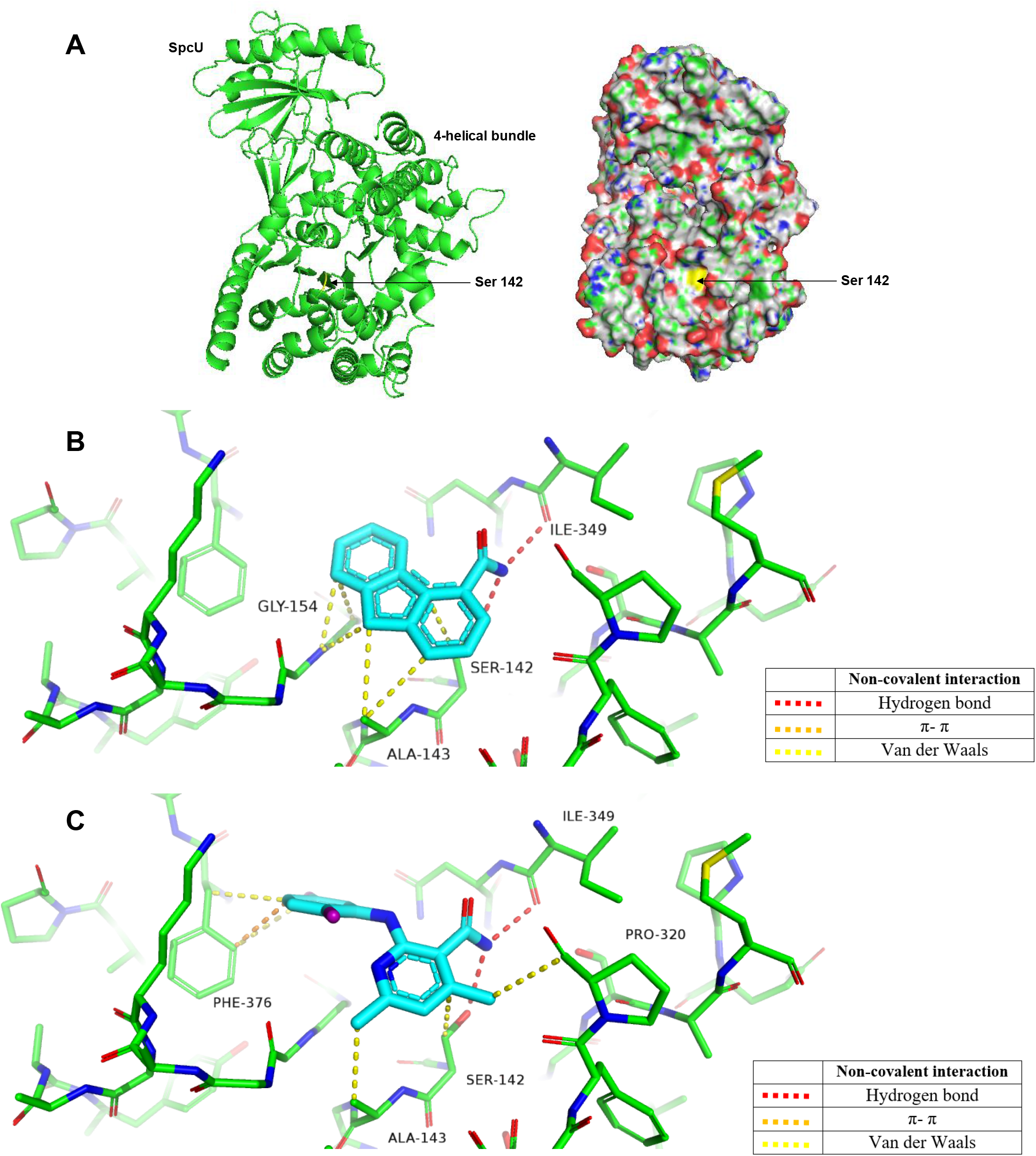
Docking poses of PSA and compound B to ExoU. (A) Structure and Connolly surface of the ExoU-SpcU complex with the catalytic serine 142 residue highlighted in yellow. Docked molecules (B) PSA and (C) compound B are rendered as sticks (carbon – cyan, nitrogen – blue, oxygen – red, chlorine – purple). Residues involved in non-covalent interactions are rendered as thin sticks (carbon – green, nitrogen – blue, oxygen – red). Non-covalent contacts are shown as dotted lines with the colour code given in the key. Non-covalent contacts analysed with ViewContacts software. Figure rendered in PyMol.

Both PSA and compound B are predicted to be bound by a large number of cooperative noncovalent interactions (Figure 6). In both cases the amide group makes a hydrogen bond interaction with Ile349 (red). Compound B has aromatic π-π stacking interactions with Phe376 (orange) and it also shows multiple van der Waals interactions with Phe376, Ala143, Ser142 and Pro320. PSA also displays multiple van der Waals interactions with Gly154, Ser142 and Ala143 with a hydrogen bond-π interaction between the amide of Gly154 and the aromatic ring of PSA.

## Discussion

### Biochemical *in vitro* analysis of ExoU with prospective small molecule inhibitors

A major challenge faced in this work was the poor yield of recombinant ExoU fusion proteins and the occurrence of similar molecular weight contaminants, which were only present in the soluble bacterial lysate after induction of ExoU expression with IPTG (Figure 1A). There is the potential that ExoU is rapidly digested by bacterial proteases shortly after induction of expression. Thus, previous purification procedures have adopted a short 3 hour induction of ExoU expression followed by immediate purification from BL21(DE3)pLysS *E. coli* [21, 28, 29, 35]. Alternate tagged variants, including glutathione-S-transferase (GST) and maltose binding protein (MBP) ExoU fusion proteins, were also quickly degraded, which was apparent from a large abundance of GST/MBP proteins with ExoU cleaved away after respective pull-downs during purification (data not shown). In this study we employed the serine protease inhibitor PMSF, which was added to C43 *E. coli* upon induction of His-tagged ExoU expression. This allowed ~5-fold greater yields of recombinant His-ExoU. The two similar molecular weight contaminating proteins seen during purification of ExoU, were identified by mass spectrometry. These were bifunctional polymyxin resistance protein ArnA and glutamine-fructose-6-phosphate aminotransferase, which have previously been documented as contaminants in IMAC purifications from *E. coli* [36] (Figure 1A). The *arnA* gene encodes a 74.3 kDa bifunctional enzyme (UDP-l-Ara4N formyltransferase/UDP-GlcA C-4”-decarboxylase), which is involved in the modification of the lipid A with 4-amino-4-deoxy-l-arabinose to confer to resistance to cationic antimicrobial peptides and antibiotics, including polymyxin [37]. ExoU localises to cellular membranes via its 4 helical bundle domain and is fully activated by eukaryotic cofactors, but mechanisms and conformational rearrangements relevant to membrane binding and catalytic activity are not yet fully understood. ExoU is toxic to *E. coli* if it is co-expressed with ubiquitin [20, 38]. Our data suggest that bifunctional polymyxin resistance protein ArnA and glutamine--fructose-6-phosphate aminotransferase could be induced in response to ExoU expression, perhaps to mitigate cell wall stress while ExoU is degraded by bacterial proteases.

### ExoU inhibitors for biochemical and mechanistic analysis

The Cayman Chemical cPLA2 Assay Kit has previously been employed to detect the phospholipase activity of recombinant ExoU [22–24]. For our analysis of ExoU phospholipase activity and to test a small panel of previously distinguished ExoU inhibitors along with certain clinical human phospholipase inhibitors, we adapted this protocol by sourcing individual reagents and making the assay compatible with a 96- and 384-well plate format. This allowed us to decrease cost and increase throughput of the assay so that it may be applicable to screening large compound libraries, which we believe should be a future focus to potentially discover novel small molecule inhibitors of ExoU. The previously distinguished ExoU inhibitors PSA, compound A and compound B (Figure 1C) exhibited low micromolar IC_50_ values for inhibition similar to those previously observed [24]. The arylsulfonamide compound, previously discovered from an independent cellular based screen [25], did not inhibit ExoU phospholipase activity *in vitro*. This does not rule out the possibility that this compound prevents ExoU mediated toxicity, by *P. aeruginosa*, through mechanisms independent of ExoU catalytic activity inhibition. As well as screening, future biochemical experiments should aim to explore the mechanisms of ExoU inhibition *in vitro* and in cells. Certain compounds may have the potential to target specific ExoU conformations or prevent interaction with activating cofactors [35] [23]. To this end, structural biological studies could be performed to aid structure guided design to improve the inhibitory activities and potency of compounds such as PSA, compound A and compound B. A co-crystal structure of ExoU with one of these compounds would be important in this endeavour. *In sillico* molecular docking simulations of prospective ExoU small molecule inhibitors to the currently solved crystal structures of the SpcU-ExoU complex, 4AKX [39] and 3TU3 [34], might give insight as to how these compounds may be optimised. Our molecular docking simulations suggest that current ExoU inhibitors (PSA, compound A and compound B) (Figure 6), bind to a solvent exposed pocket in the catalytic domain, forming various polar interactions.

### Inhibition of ExoU expressed in transfected HeLa cells

Transfected mammalian cells undergo rapid cellular lysis when induced to express WT, but not S142A ExoU [27]. In the HeLa transfection model, PSA, compound A and compound B, but not arylsulfonamide 1, were able to mitigate cytotoxicity induced after ExoU expression. The effects observed with PSA, compound A and compound B were dose responsive and statistically similar. The data suggest that ExoU is degraded by cellular proteases, as ExoU stability was increased in the presence of PMSF but not MG132 (supplementary Figure 3). It was previously observed that ExoU becomes ubiquitinated in mammalian cells at Lys178 [33, 39]. This modification does not seem to influence the toxicity exerted by WT ExoU phospholipase activity [39]. Ubiquitinated S142A ExoU was found to be targeted to acidic organelles [39], which might contribute to its turnover by endosomal proteases. In our study, ExoU protein abundance did not increase in the presence of the proteasome inhibitor MG132, but did in the presence of the protease inhibitor PMSF. The inositol polyphosphate phosphatase SopB is a *Salmonella* type III effector becomes ubiquitinated at Lys6 after delivery to mammalian cells [40]. This modification also does not affect SopB stability or membrane association, but was found to extend temporal association with *Salmonella*-containing vacuoles (SCVs) [40]. It is yet to be fully explored whether or not ubiquitination of ExoU contributes to activation or molecular rearrangement, but there is evidence to suggest that ubiquitination of ExoU promotes endosomal association [39], which perhaps serves as a defence mechanism against ExoU mediated cytotoxicity.

S142A ExoU could be detected more readily in cellular lysates [33], perhaps due to the fact that WT ExoU expression was acutely lethal and thus could not accumulate to an abundance that was reasonably detectable in transfected HeLa cells (Figure 2A). Therefore, by inducing expression of S142A ExoU, in the presence of ExoU inhibitors, we observed that compounds A and B, but not PSA or arylsulfonamide 1, caused a reduction in the total amount of S142A ExoU (Figure 3). PSA, compound A and compound B exerted similar IC_50_ values for ExoU inhibition *in vitro* (Figure 1B) and had similar protective effects in WT ExoU expressing HeLa cells (Figure 2B, C, D and E). All three compounds possess the ability inhibit ExoU in HeLa cells but compounds A and B may also potentiate degradation of ExoU by endogenous proteases.

### A scratch and infection assay to evaluate the therapeutic potential of ExoU inhibitors

Corneal cells of the human eye form a non-polar apical barrier, which is important for function and protection against infection [41]. Infections usually arise after physical damage to the cornea by opportunistic pathogens [41, 42]. We found that fully confluent HCE-T cells were not susceptible to infection by PA103 (Figure 4A), but sub-confluent cells (~90%) were. By applying a scratch to the bottom of the well of fully confluent monolayer of HCE-T cells, we could simulate injury and allow T3SS and ExoU mediated cytotoxicity from PA103, which propagated over time from the periphery of the scratch, extending outwards. This movement of infection, spreading from the border of the scratch and towards neighbouring cells, may offer insights into the mechanisms of infection and disease progression *in vivo*. Not only could we simulate injury in this way, the scratch had the advantage of providing a point of focus from which to observe the effects of ExoU mediated cytotoxicity and potential protective effects of antibiotics and prospective ExoU small molecule inhibitors. This was essential for quantification of the effects of inhibitors. In the scratch assay, PSA, compound A and compound B were effective in the low micromolar region, with significant protection of HCE-T cells from PA103 infection by compound B at 0.5 μM (Figure 4C). The toxicity analysis in HCE-T cells indicated that all these compounds were well tolerated in the high micromolar concentrations (>30 μM) (Figure 4B), suggesting that with topical administration, ExoU inhibitors might achieve effective inhibitory concentrations.

The scratch assay was extendable by employing moxifloxacin, a fluoroquinolone commonly used to treat eye infections that inhibits bacterial DNA synthesis [8], at an established MIC to manage bacterial load, whilst still maintaining the effects of ExoU mediated cytotoxicity (Figure 5B). We believe that this assay could be employed as a useful tool to analyse the effects of T3SS cytotoxicity in cell culture models for longer times. Using moxifloxacin to manage bacterial load in scratch and infection assays may offer insights into mechanisms of disease progression and how infections could respond to extended treatments. Only when used in combination with moxifloxacin were PSA, compounds A and compound B effective at protecting HCE-T cells from infection over 24 hours (Figure 5D). This is likely due to the bacteria, when incubated with HCE-T cells for 24 hours, growing exponentially leading to cytotoxic effects independent of ExoU expression [43]. In the instance of combinational treatment of compound B (10 μM) and moxifloacin (at the MIC), partial wound closure was observed (Figure 5E). With 10 μM compound A and to a lesser extent with 10 μM of PSA, with moxifloxacin, narrowing of the scratch was observed. The data, therefore, suggest that compounds A and B could be more effective than PSA at protecting HCE-T cells from ExoU for longer treatment courses. The stability of S142A ExoU was only significantly decreased by compounds A and B but not PSA, in HeLa cells (Figure 3), despite the three compounds possessing similar IC_50_ values for *in vitro* inhibition of ExoU phospholipase activity (Figure 1C). An explanation for the difference may be due to better cellular penetration of compounds A and B compared to PSA.

An interesting finding, from our scratch and infection assay, was that tobramycin, applied at the established MIC, completely abrogated any observable effect of ExoU mediated cytotoxicity (Figure 5A, B and C). Despite the fact there were more PA013 colony forming units detected in the HCE-T cell medium of tobramycin exposed cells than ones exposed to moxifloxacin (Figure 5A), complete wound healing and no cellular lysis was observed only for tobramycin treated HCE-T cells (Figure 5 B and C). Tobramycin is an aminoglycoside, which inhibits protein synthesis by binding to bacterial 30S and 50S ribosomes [44]. Tobramycin did not inhibit the *in vitro* phospholipase activity of ExoU (supplementary Figure 6), which would suggest that abolished toxicity induced by PA103, in the presence of tobramycin at the MIC, was due to inhibited translocation of ExoU into HCE-T cells. Perhaps, in response to tobramycin, the PA103 did not prioritise synthesis of T3SS proteins or altered T3SS transcriptional regulation lead to abolished ExoU mediated cytotoxicity [45]. A previous study has indicated that at lower concentrations of tobramycin, lethality was facilitated through protein synthesis inhibition (<4 μg/ml), but at high concentrations (≥8 μg/ml) *P. aeruginosa* outer membranes were shown to deteriorate [44]. Future studies could aim to monitor expression of the four key T3SS transcription factors (ExsA, ExsC, ExsD and ExsE) [45] or T3SS associated proteins [46], after exposure of *P. aeruginosa* to tobramycin, using immunoblotting and Q-PCR. Such data could inform on whether or not tobramycin, or similar aminoglycosides, could be considered as a viable treatment approaches for microbial keratitis, where cytotoxicity is driven by the T3SS of *P. aeruginosa*.

Our analysis suggests that pharmacological targeting of ExoU may be compatible with antibiotic usage, whereby inhibitors of ExoU serve as an adjuvant therapy. We believe that there is potential for *ex vivo* and *in vivo* analysis where ExoU inhibitors could be employed, alone or as an adjuvant in combination with antibiotics. Indeed, these ExoU inhibitors were not bactericidal, therefore, in a therapeutic context, inhibitors would serve to mitigate the rapid toxicity induced by ExoU, while antibiotics clear the infection.

## Acknowledgments

We would like to personally thank Professor Dara Frank for kindly providing us with PA103 and PA103 mutant strains used in this study. We would also like to thank the University of Liverpool Centre for Proteome research (CPR) for use of the mass spectrometry suite.

## Supplementary figures

**Supplementary Figure 1:**
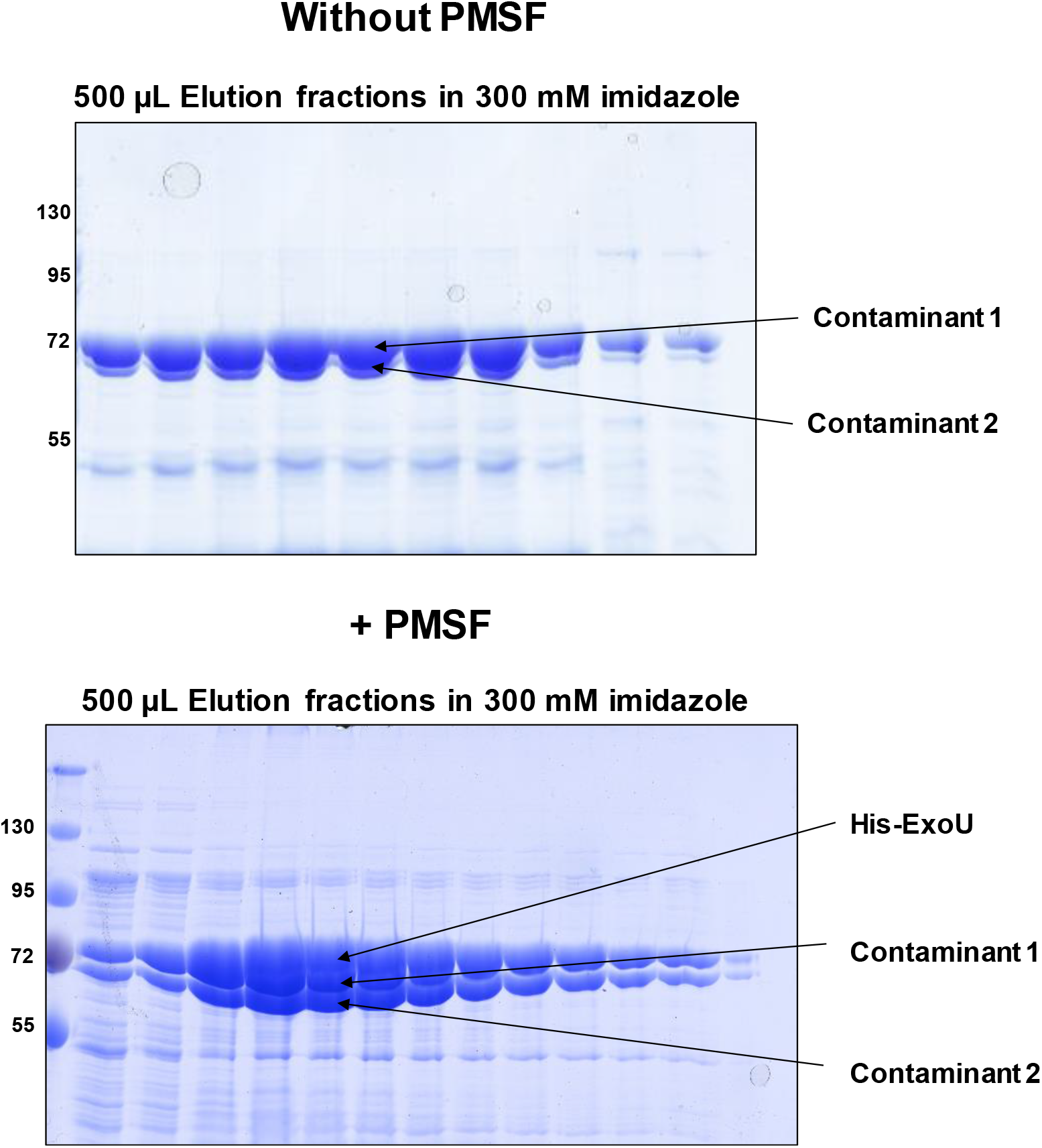
Expression of His-tagged ExoU in the presence of PMSF. C43(DE3) E. coli were grown to 0.8 OD_600_ followed by the addition of IPTG and 100 μM PMSF. ExoU was expressed for 3 hours with the addition of 100 μM PMSF at each hour before centrifugation of E. coli and purification by immobilised metal affinity chromatography (IMAC).

**Supplementary Figure 2:**
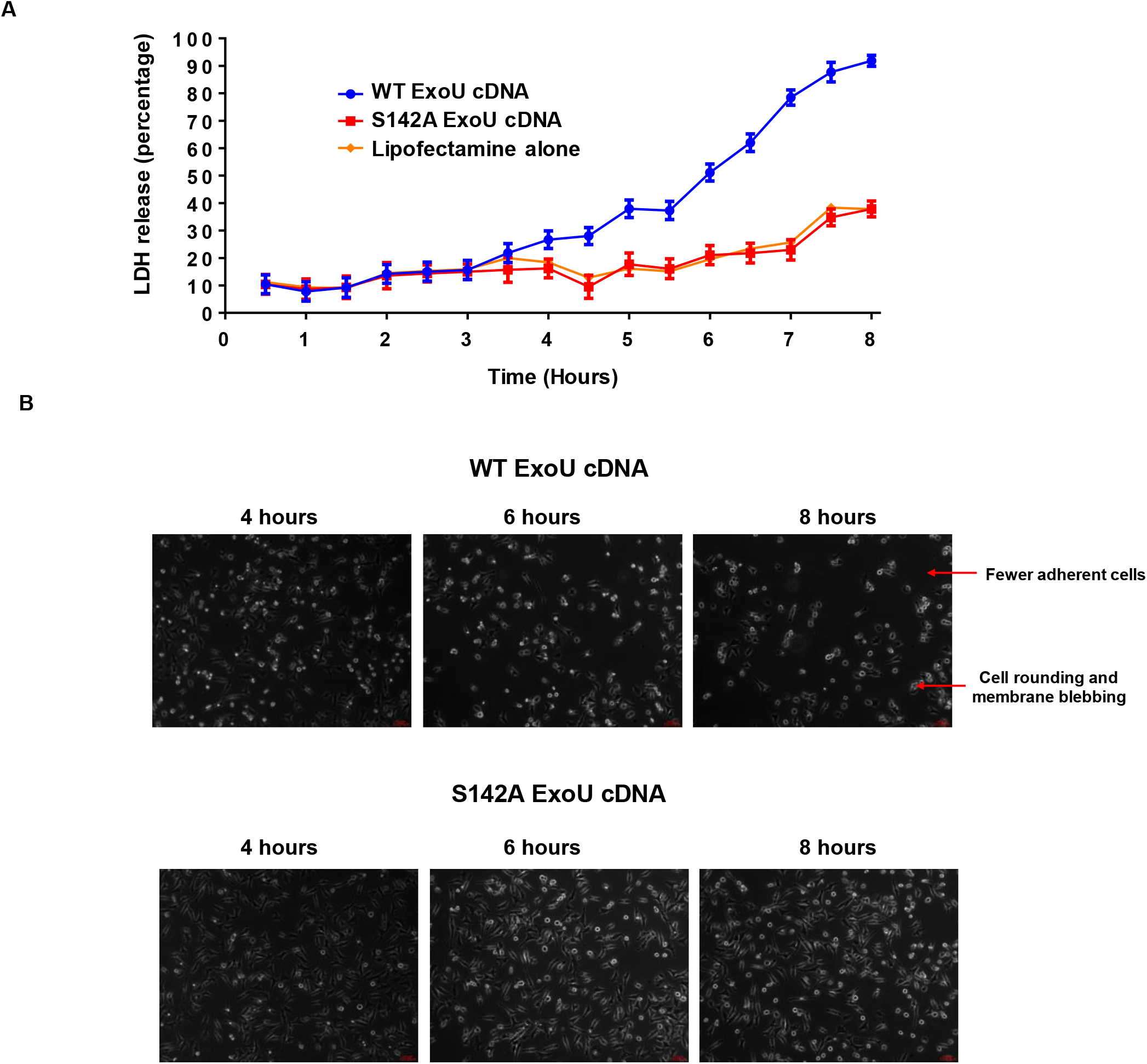
Effect of WT and S142A ExoU expression on HeLa cell viability. HeLa cells were seeded in wells of a 6-well plate for 24 hours. Lipofectamine was used to transfect pcDNA5/FRT/TO encoding WT or S142A ExoU cDNA (1 μg per well) for 12 hours, followed by the addition of 10 μg/ml of tetracycline to induction of ExoU expression. (A) LDH assay time course analysis of HeLa cells transfected to express WT ExoU (blue) or S142A ExoU (red). (B) Brightfield images taken of transfected HeLa cells after induced expression of either WT or S142A ExoU at the indicated time points.

**Supplementary Figure 2:**
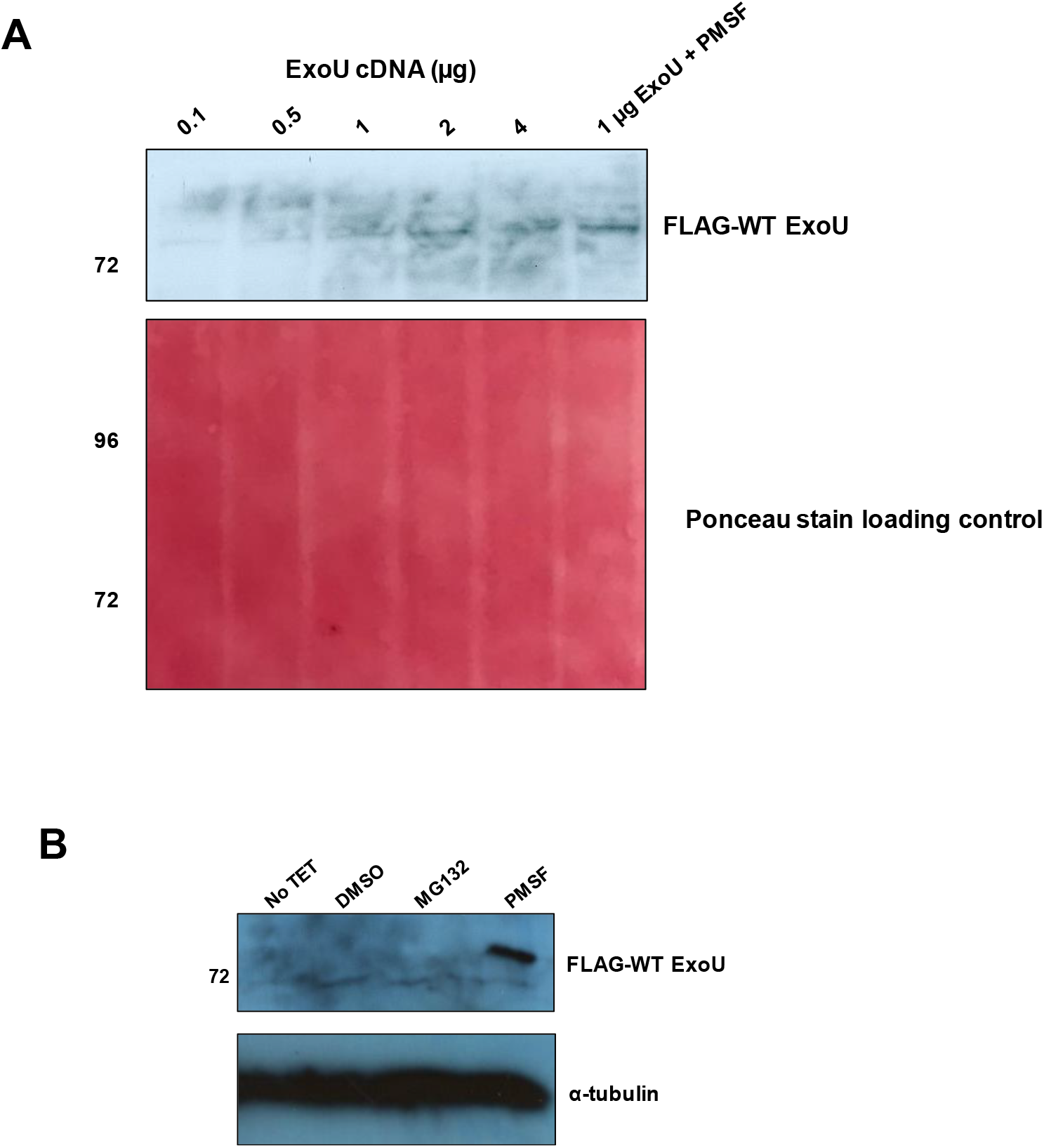
Effect of WT and S142A ExoU expression on HeLa cell viability. (A) HeLa cells were seeded in wells of a 6-well plate for 24 hours. Lipofectamine was used to transfect varying amounts of pcDNA5/FRT/TO encoding WT ExoU cDNA for 12 hours, followed by the addition of 10 μg/ml of tetracycline to induction of ExoU expression. Western blot analysis reveals the total abundance of FLAG-tagged WT-ExoU. (B) Western blot analysis of FLAG-tagged WT ExoU expression in TET induced HeLa cells after incubation with 50 μM MG132 or PMSF for 6 hours.

**Supplementary Figure 4:**
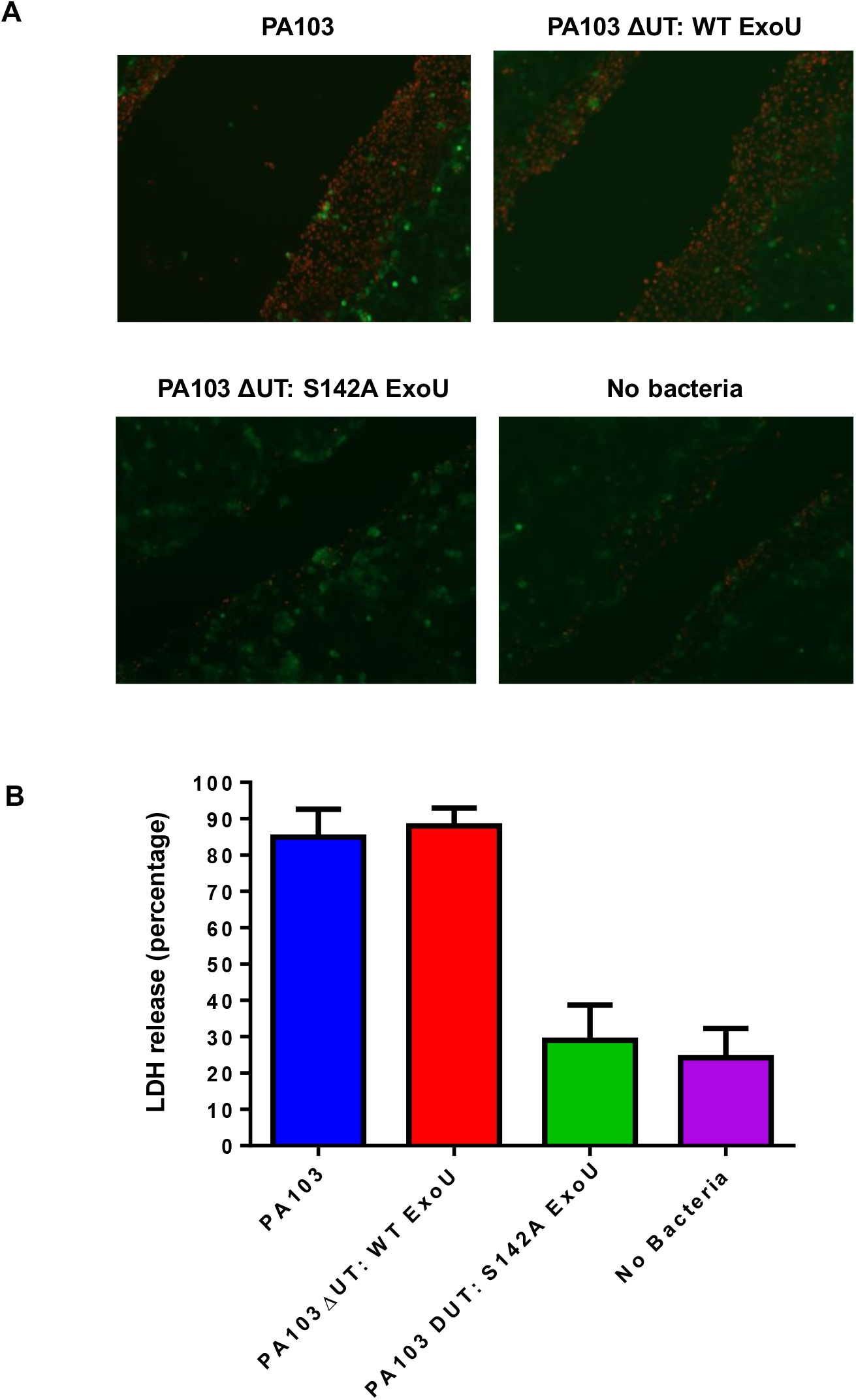
Infection of scratched HCE-T cells with PA103 mutants. HCE-T were gown to full confluence and a scratch applied to the well bottom prior to infection with either PA103, PA103 ΔUT: ExoU or PA103 ΔUT: S142A ExoU at an MOI of 2.5 for 6 hours, followed by analysis by Live/Dead fluorescence microscopy (A) or LDH release (B).

**Supplementary Figure 5:**
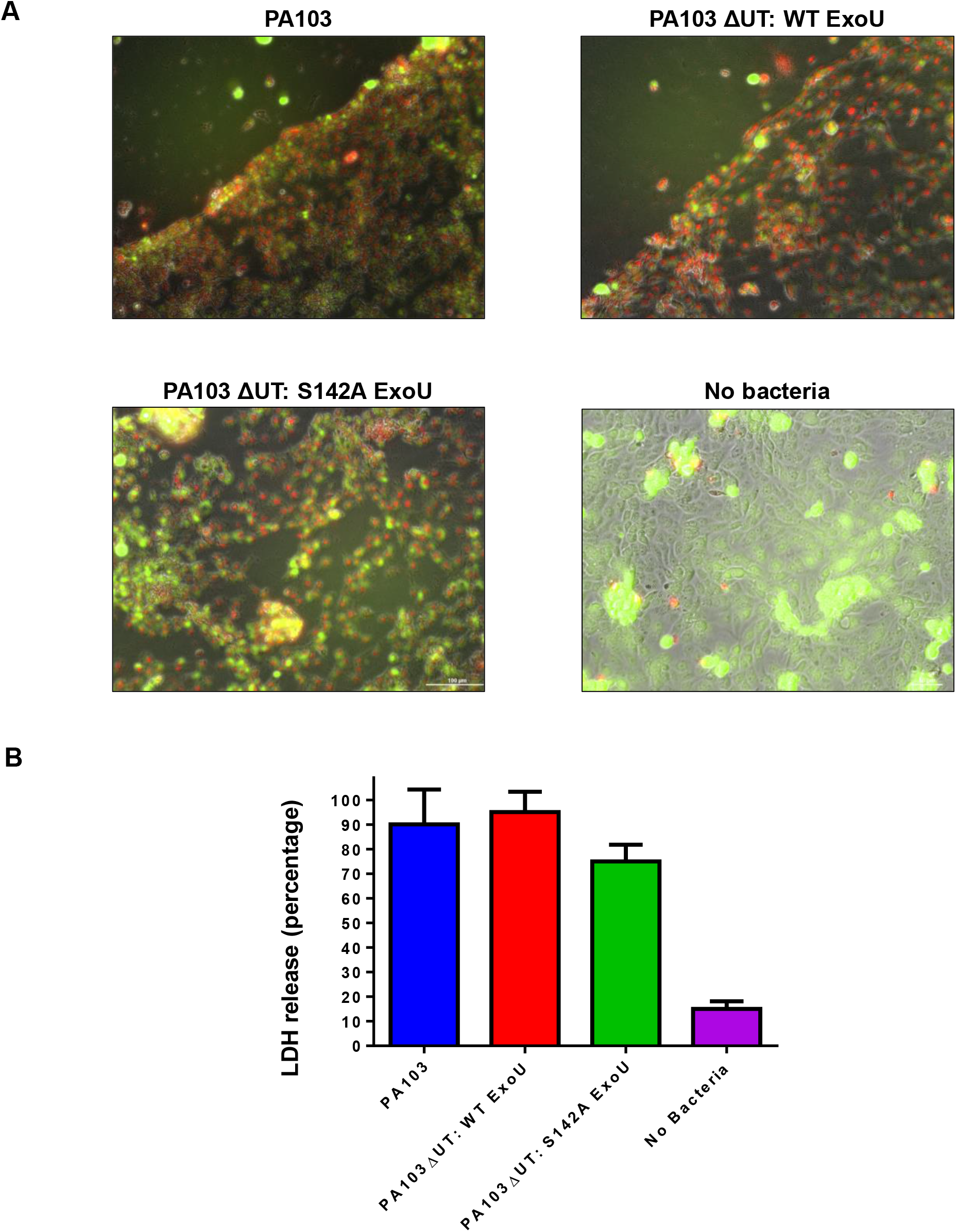
Toxicity of PA103 mutants to scratched HCE-T cells after 24 hours incubation without antibiotic present. HCE-T were gown to full confluence and a scratch applied to the well bottom prior to infection with either PA103, PA103 ΔUT: ExoU or PA103 ΔUT: S142A ExoU at an MOI of 2.5 for 24 hours, followed by analysis by Live/Dead fluorescence microscopy (A) or LDH release (B).

**Supplementary Figure 6:**
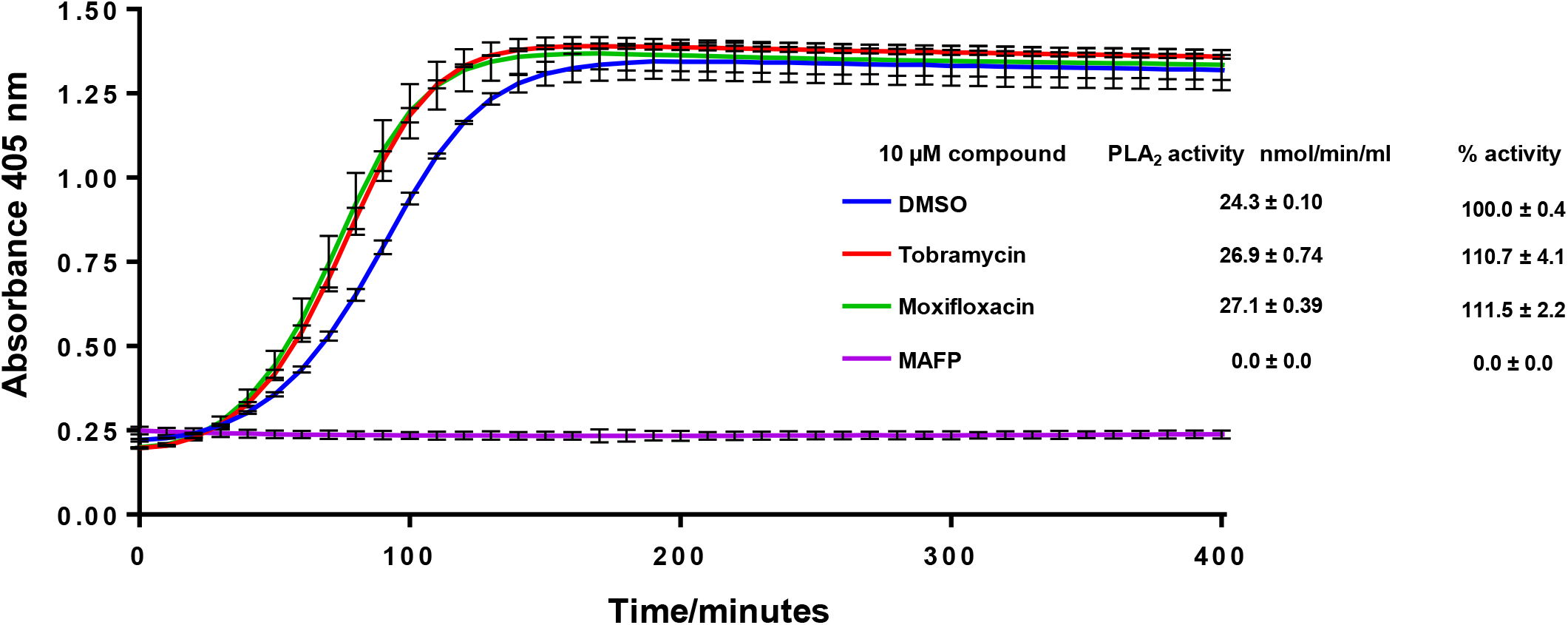
Moxifloxacin and tobramycin do not inhibit recombinant ExoU. The hydrolysis of arachidonoyl Thio-PC substrate by ExoU was assessed in the presence of 10 μM of moxifloxacin or tobramycin. To each reaction, ubiquitin and PIP2 were added in order to allow induction of ExoU phospholipase activity.

## References

1. Stewart, R.M., et al., Genetic characterization indicates that a specific subpopulation of Pseudomonas aeruginosa is associated with keratitis infections. J Clin Microbiol, 2011. 49(3): p. 993–1003.

2. Newman, J.W., R.V. Floyd, and J.L. Fothergill, The contribution of Pseudomonas aeruginosa virulence factors and host factors in the establishment of urinary tract infections. FEMS Microbiol Lett, 2017. 364(15).

3. Schulert, G.S., et al., Secretion of the toxin ExoU is a marker for highly virulent Pseudomonas aeruginosa isolates obtained from patients with hospital-acquired pneumonia. J Infect Dis, 2003. 188(11): p. 1695–706.

4. Roy-Burman, A., et al., Type III protein secretion is associated with death in lower respiratory and systemic Pseudomonas aeruginosa infections. J Infect Dis, 2001. 183(12): p. 1767–74.

5. Hauser, A.R., et al., Type III protein secretion is associated with poor clinical outcomes in patients with ventilator-associated pneumonia caused by Pseudomonas aeruginosa. Crit Care Med, 2002. 30(3): p. 521–8.

6. Migiyama, Y., et al., Pseudomonas aeruginosa Bacteremia among Immunocompetent and Immunocompromised Patients: Relation to Initial Antibiotic Therapy and Survival. Jpn J Infect Dis, 2016. 69(2): p. 91–6.

7. Tacconelli, E., et al., Discovery, research, and development of new antibiotics: the WHO priority list of antibiotic-resistant bacteria and tuberculosis. The Lancet Infectious Diseases, 2018. 18(3): p. 318–327.

8. Al-Mujaini, A., et al., Bacterial keratitis: perspective on epidemiology, clinico-pathogenesis, diagnosis and treatment. Sultan Qaboos Univ Med J, 2009. 9(2): p. 184–95.

9. Lombardi, C., et al., Structural and Functional Characterization of the Type Three Secretion System (T3SS) Needle of Pseudomonas aeruginosa. Front Microbiol, 2019. 10: p. 573.

10. Galle, M., I. Carpentier, and R. Beyaert, Structure and function of the Type III secretion system of Pseudomonas aeruginosa. Curr Protein Pept Sci, 2012. 13(8): p. 831–42.

11. Finck-Barbancon, V., et al., ExoU expression by Pseudomonas aeruginosa correlates with acute cytotoxicity and epithelial injury. Mol Microbiol, 1997. 25(3): p. 547–57.

12. Fernandez-Barat, L., et al., Intensive care unit-acquired pneumonia due to Pseudomonas aeruginosa with and without multidrug resistance. J Infect, 2017. 74(2): p. 142–152.

13. Sato, H. and D.W. Frank, ExoU is a potent intracellular phospholipase. Mol Microbiol, 2004. 53(5): p. 1279–90.

14. Foulkes, D.M., et al., Pseudomonas aeruginosa Toxin ExoU as a Therapeutic Target in the Treatment of Bacterial Infections. Microorganisms, 2019. 7(12).

15. Sato, H., et al., The mechanism of action of the Pseudomonas aeruginosa-encoded type III cytotoxin, ExoU. EMBO J, 2003. 22(12): p. 2959–69.

16. Phillips, R.M., et al., In vivo phospholipase activity of the Pseudomonas aeruginosa cytotoxin ExoU and protection of mammalian cells with phospholipase A2 inhibitors. J Biol Chem, 2003. 278(42): p. 41326–32.

17. Rabin, S.D. and A.R. Hauser, Functional regions of the Pseudomonas aeruginosa cytotoxin ExoU. Infect Immun, 2005. 73(1): p. 573–82.

18. Tessmer, M.H., et al., Cooperative Substrate-Cofactor Interactions and Membrane Localization of the Bacterial Phospholipase A2 (PLA2) Enzyme, ExoU. J Biol Chem, 2017. 292(8): p. 3411–3419.

19. Tessmer, M.H., et al., Identification of a ubiquitin-binding interface using Rosetta and DEER. Proc Natl Acad Sci U S A, 2018. 115(3): p. 525–530.

20. Anderson, D.M., et al., Ubiquitin and ubiquitin-modified proteins activate the Pseudomonas aeruginosa T3SS cytotoxin, ExoU. Mol Microbiol, 2011. 82(6): p. 1454–67.

21. Zhang, A., J.L. Veesenmeyer, and A.R. Hauser, Phosphatidylinositol 4,5-Bisphosphate-Dependent Oligomerization of the Pseudomonas aeruginosa Cytotoxin ExoU. Infect Immun, 2018. 86(1).

22. Tyson, G.H., et al., A novel phosphatidylinositol 4,5-bisphosphate binding domain mediates plasma membrane localization of ExoU and other patatin-like phospholipases. J Biol Chem, 2015. 290(5): p. 2919–37.

23. Tyson, G.H. and A.R. Hauser, Phosphatidylinositol 4,5-bisphosphate is a novel coactivator of the Pseudomonas aeruginosa cytotoxin ExoU. Infect Immun, 2013. 81(8): p. 2873–81.

24. Lee, V.T., et al., Pseudolipasin A is a specific inhibitor for phospholipase A2 activity of Pseudomonas aeruginosa cytotoxin ExoU. Infect Immun, 2007. 75(3): p. 1089–98.

25. Kim, D., et al., Identification of arylsulfonamides as ExoU inhibitors. Bioorg Med Chem Lett, 2014. 24(16): p. 3823–5.

26. Tam, C., et al., Mutation of the phospholipase catalytic domain of the Pseudomonas aeruginosa cytotoxin ExoU abolishes colonization promoting activity and reduces corneal disease severity. Exp Eye Res, 2007. 85(6): p. 799–805.

27. Sato, H. and D.W. Frank, Intoxication of host cells by the T3SS phospholipase ExoU: PI(4,5)P2-associated, cytoskeletal collapse and late phase membrane blebbing. PLoS One, 2014. 9(7): p. e103127.

28. Schmalzer, K.M., M.A. Benson, and D.W. Frank, Activation of ExoU phospholipase activity requires specific C-terminal regions. J Bacteriol, 2010. 192(7): p. 1801–12.

29. Benson, M.A., et al., Induced conformational changes in the activation of the Pseudomonas aeruginosa type III toxin, ExoU. Biophys J, 2011. 100(5): p. 1335–43.

30. Jones, G., et al., Development and validation of a genetic algorithm for flexible docking. J Mol Biol, 1997. 267(3): p. 727–48.

31. Kuhn, B., et al., Rationalizing tight ligand binding through cooperative interaction networks. J Chem Inf Model, 2011. 51(12): p. 3180–98.

32. Benson, M.A., K.M. Schmalzer, and D.W. Frank, A sensitive fluorescence-based assay for the detection of ExoU-mediated PLA(2) activity. Clin Chim Acta, 2010. 411(3-4): p. 190–7.

33. Stirling, F.R., et al., Eukaryotic localization, activation and ubiquitinylation of a bacterial type III secreted toxin. Cell Microbiol, 2006. 8(8): p. 1294–309.

34. Halavaty, A.S., et al., Structure of the type III secretion effector protein ExoU in complex with its chaperone SpcU. PLoS One, 2012. 7(11): p. e49388.

35. Anderson, D.M., et al., Identification of the major ubiquitin-binding domain of the Pseudomonas aeruginosa ExoU A2 phospholipase. J Biol Chem, 2013. 288(37): p. 26741–52.

36. Robichon, C., et al., Engineering Escherichia coli BL21(DE3) derivative strains to minimize E. coli protein contamination after purification by immobilized metal affinity chromatography. Appl Environ Microbiol, 2011. 77(13): p. 4634–46.

37. Breazeale, S.D., A.A. Ribeiro, and C.R. Raetz, Oxidative decarboxylation of UDP-glucuronic acid in extracts of polymyxin-resistant Escherichia coli. Origin of lipid a species modified with 4-amino-4-deoxy-L-arabinose. J Biol Chem, 2002. 277(4): p. 2886–96.

38. Anderson, D.M., et al., Ubiquitin activates patatin-like phospholipases from multiple bacterial species. J Bacteriol, 2015. 197(3): p. 529–41.

39. Gendrin, C., et al., Structural basis of cytotoxicity mediated by the type III secretion toxin ExoU from Pseudomonas aeruginosa. PLoS Pathog, 2012. 8(4): p. e1002637.

40. Knodler, L.A., et al., Ubiquitination of the bacterial inositol phosphatase, SopB, regulates its biological activity at the plasma membrane. Cell Microbiol, 2009. 11(11): p. 1652–70.

41. Bolanos-Jimenez, R., et al., Ocular Surface as Barrier of Innate Immunity. Open Ophthalmol J, 2015. 9: p. 49–55.

42. Lim, B.X., V.T.C. Koh, and M. Ray, Microbial characteristics of post-traumatic infective keratitis. Eur J Ophthalmol, 2018. 28(1): p. 13–18.

43. Ongradi, J., G. Kulcsar, and L. Bertok, Toxicity of microbial products in cell culture. Folia Microbiol (Praha), 1984. 29(6): p. 450–4.

44. Bulitta, J.B., et al., Two mechanisms of killing of Pseudomonas aeruginosa by tobramycin assessed at multiple inocula via mechanism-based modeling. Antimicrob Agents Chemother, 2015. 59(4): p. 2315–27.

45. Williams McMackin, E.A., et al., Fitting Pieces into the Puzzle of Pseudomonas aeruginosa Type III Secretion System Gene Expression. J Bacteriol, 2019. 201(13).

46. Galan, J.E., et al., Bacterial type III secretion systems: specialized nanomachines for protein delivery into target cells. Annu Rev Microbiol, 2014. 68: p. 415–38.

